# Transcranial magnetic stimulation entrains alpha oscillatory activity in occipital cortex

**DOI:** 10.1101/2021.01.09.426064

**Authors:** Yong-Jun Lin, Lavanya Shukla, Laura Dugué, Antoni Valero-Cabré, Marisa Carrasco

## Abstract

Parieto-occipital alpha rhythms (8-12 Hz) underlie cortical excitability and influence visual performance. Whether the synchrony of intrinsic alpha rhythms in the occipital cortex can be entrained by transcranial magnetic stimulation (TMS) is an open question. We applied 4-pulse, 10-Hz rhythmic TMS to entrain intrinsic alpha oscillators targeting right V1/V2, and tested four predictions with concurrent electroencephalogram (EEG): (1) progressive enhancement of entrainment across time windows, (2) output frequency specificity, (3) dependence on the intrinsic oscillation phase, and (4) input frequency specificity to individual alpha frequency (IAF) in the neural signatures. Two control conditions with an equal number of pulses and duration were arrhythmic-active and rhythmic-sham stimulation. The results confirmed the first three predictions. Rhythmic TMS bursts significantly entrained local neural activity. Near the stimulation site, evoked oscillation amplitude and inter-trial phase coherence (ITPC) were increased for 2 and 3 cycles, respectively, after the last TMS pulse. Critically, ITPC following entrainment positively correlated with IAF rather than with the degree of similarity between IAF and the input frequency (10 Hz). Thus, we entrained alpha-band activity in occipital cortex for ~3 cycles (~300 ms), and IAF predicts the strength of entrained occipital alpha phase synchrony indexed by ITPC.

---

Brain oscillations play an important functional role in perception, attention and cognition^1–3^. Specifically, alpha-band (8-12 Hz) neural oscillations reflect ongoing sensory processing. Alpha power inversely correlates with excitability in vision^4–9^, audition^10^, and somatosensation^11^. In the visual cortex, the parieto-occipital alpha-band phase correlates with baseline cortical excitability (phosphene threshold)^12^, gamma-band (40-100 Hz) power^13^, and spiking responses^14^. In V1, alpha and gamma rhythms index feedback and feedforward processing, respectively^15,16^. Theories of oscillations postulate that alpha power reflects an active inhibition of task-irrelevant sensory signals^17–19^ whereas alpha phase reflects pulsed inhibition^17,20^, cyclic perceptual sampling^2^ or temporal expectation^21^. Here we aimed to use transcranial magnetic stimulation (TMS) in each trial to directly manipulate local alpha rhythms while we assess its neurophysiological impact with concurrent EEG.

The topography of alpha power is altered by spatial attention. Typically, in endogenous (voluntary) spatial attention tasks, alpha power contralateral to the cued visual field shows reduced alpha power while the ipsilateral side shows increased alpha power^22–29^. Moreover, this lateralized alpha power modulation could reach V1^29,30^. The parieto-occipital alpha-band activity is linked with the allocation of covert spatial attention and receives feedback control from frontoparietal cortices. The disruption of anticipatory pre-stimulus alpha rhythms on the occipital cortex brought about by transcranial magnetic stimulation (TMS) on the right intraparietal sulcus (IPS) or right frontal eye field (FEF) has been associated with deteriorated visual performance^31–33^. With Granger causality analyses applied to magnetoencephalogram data, alpha power modulation in the right FEF has been reported to predict alpha activity in V1, indicating feedback control^34^. Thus, V1/V2 may receive feedback control from FEF and IPS, two crucial nodes of the dorsal attention network.

Neural entrainment –the progressive phase alignment of intrinsic oscillators by external sources of rhythmic stimulation (reviews^35–37^)– modulates behavioral performance in many visual^38–44^, tactile^45^, motor^46^, and memory^47^ tasks. Behavioral studies with concurrent TMS-EEG can shed light on neural oscillatory processes of behavior^43,44,46–48^. Entrainment from higher-order to sensory areas has been proposed to underlie top-down modulations of attention^40,42,43,49–52^. Although visual signals in visuo-occipital areas are modulated by attention^53–56^, a fundamental question is whether and how the occipital cortex would respond to entrainment while participants perform a visual task, even without attention being explicitly manipulated. To address this question, we utilized a short-burst rhythmic TMS protocol along with concurrent EEG recordings^43,57^, targeting V1/V2 with visual field mapping by fMRI and/or phosphene induction.

For the frequency of alpha-band stimulation, previous short-burst TMS studies used either 10 Hz^38,48^ or individual alpha frequency (IAF) as the entrainment frequency^57,58^. IAF is a stable neurophysiological trait marker at rest^59^, which increases with task demand^60,61^ and reflects the task-required temporal integration window^62^. However, the assumption that matching the entraining frequency with IAF is important has not been directly tested in concurrent TMS-EEG studies^37^. Our strategy was to use 10 Hz as the entraining frequency and examine whether the degree of similarity between IAF and 10 Hz predicted the magnitude of phase-locked activities following entrainment.

Here we asked how the occipital cortex responds to entrainment while controlling spatial attention allocation to be distributed across the left and right visual fields, where the stimuli appeared. We aimed to entrain alpha-band synchrony in early visual areas with rhythmic 10 Hz TMS and examine the neural signatures of entrainment with EEG. We hypothesized that the occipital alpha-band activity can be entrained by targeting V1/V2 because intrinsic alpha oscillators exist in V1^14,63^ and V2^64^. Given that V1/V2 play a pivotal role in early visual information processing, it is important to find out whether entrainment with rhythmic TMS can be established in these areas and how long such entrainment would last. We adopted arrhythmic-active TMS and rhythmic-sham stimulation control conditions, similar to previous TMS-EEG entrainment studies^43,57^. We systematically analyzed the temporal dynamics in multiple time windows during and after stimulation bursts. Although modulations of neural activity do not necessarily lead to changes in behavior, our secondary goal was to examine whether entrained alpha-band oscillations targeting early visual cortex (V1/V2) could affect perceptual discriminability and/or criteria in a visual discrimination task.

Based on our hypothesis that occipital alpha-band activity can be entrained by targeting V1/V2 with 10 Hz TMS bursts, we tested four specific predictions characterizing TMS-driven entrainment (the first three have been previously tested in frontal and parietal locations^43,57^): (1) *Progressive enhancement of entrainment*: the strength of phase-locked activity, as measured by evoked (phase-locked) oscillation amplitude (as opposed to induced, non-phase-aligned, oscillation amplitude^65^) and ITPC (consistency of phase alignment)^43,57^, should be enhanced only after the second TMS pulse of the 4-pulse burst, because the alpha periodicity of the entraining rhythm is not defined until then. Otherwise, the results could simply reflect phase-resetting due to the first TMS pulse. (2) *Output frequency specificity*: entrained phase-locked activity should peak around the 10-Hz TMS frequency. (3) *Dependence on the intrinsic oscillation phase:* entrained activity should depend on the pre-TMS alpha phase^57^. Were the enhanced phase-locked activity due to reverberation of the imposed rhythm, rather than entrainment of intrinsic oscillators, the entrained activity should be similar regardless of pre-TMS alpha phase. (4) *Input frequency specificity*: The strength of the entrained activity should correlate with the degree of similarity between IAF and 10 Hz, given that IAF may be the intrinsic oscillation frequency of occipital alpha oscillators^57^.

## Materials and Methods

### Participants

All 11 participants (6 male, 5 female; 9 naïve to the purpose of the study; 20-47 years; M=29.9; SD=6.8) provided written informed consent. One participant’s data were excluded due to excessive involuntary blinking. The number of participants was determined based on a power analysis (β > 0.8 at alpha = 0.05) from bootstrapping 5 participants’ data, and was comparable to previous alpha-band entrainment TMS-EEG studies in the parietal cortex^48,57^. NYU institutional review board approved the protocol (IRB #i14-00788), which followed the Declaration of Helsinki and safety guidelines for TMS experiments^66^.

### Apparatus

The stimuli were presented on a ViewSonic P220f monitor. The screen resolution was 800(H) × 600(V) at 120Hz. The viewing distance was 57 cm, set by a chin-rest. The stimuli presentation code was written in MATLAB with Psychtoolbox 3^67,68^. To linearize stimuli contrast, the monitor’s gamma function was measured with a ColorCAL MKII Colorimeter (Cambridge Research Systems).

The TMS pulses were delivered with a 70-mm figure-of-eight coil controlled by a Magstim Super Rapid^2^ Plus^1^ system. The coil, whose handle pointed rightward with respect to the midline of the skull, was supported by a mechanical arm and hand-held tangentially to the skull; its positioning was always guided by a Brainsight neuronavigation system (Rogue Research) with ~1 mm precision. Two identical TMS coils were alternated every 2 blocks of testing to prevent overheating. Infrared reflecting markers were attached to the TMS coil and the participant’s head to extract the relative 3D positions in real time; they were displayed on each participant’s MRI structural scan.

The EEG system consisted of a Brain Products actiCHamp amplifier and TMS-compatible Easycap actiCAP slim caps. The EEG cap layout, following the international 10-20 system^69^, included a grid of 63 TMS compatible 6mm-thick electrodes, including a ground electrode placed at Fpz and a reference electrode placed at FCz. Impedance of all electrodes after conducting gel application was kept below 25k Ohm during the whole experimental session. The EEG recording software was BrainVision Recorder. An in-house built TMS triggering and EEG event registering device with at least 0.4 ms resolution ensured timing precision at 2500 Hz EEG sampling rate^70^ (**Fig. S1A**).

### Visual discrimination task

With the goal of facilitating entrainment, we ensured that participants were engaged in a visual task. The participants performed an orientation discrimination task. Each trial began with a fixation period, followed by a 25 ms neutral pre-cue indicating that the target was equally likely to appear in the lower left or lower right visual field (**Fig. 1A**). After a 500 ms inter-stimulus interval (ISI), two Gabor patches and a response cue, indicating which patch was the target, appeared simultaneously. The Gabor patches (achromatic; 4 cpd; σ 0.42°) lasted 50 ms. Participants were asked to indicate whether the target patch was slightly clockwise or counterclockwise relative to vertical via a key press (right index finger for the ‘/’ key; left index finger for the ‘z’ key). A high pitch (700 Hz) signaled a correct response and a low tone (400 Hz) signaled an incorrect response. Only after the participants responded, the next trial would begin. The inter-trial interval was randomly chosen among 2500, 2750, or 3000 ms.

**Figure 1.**
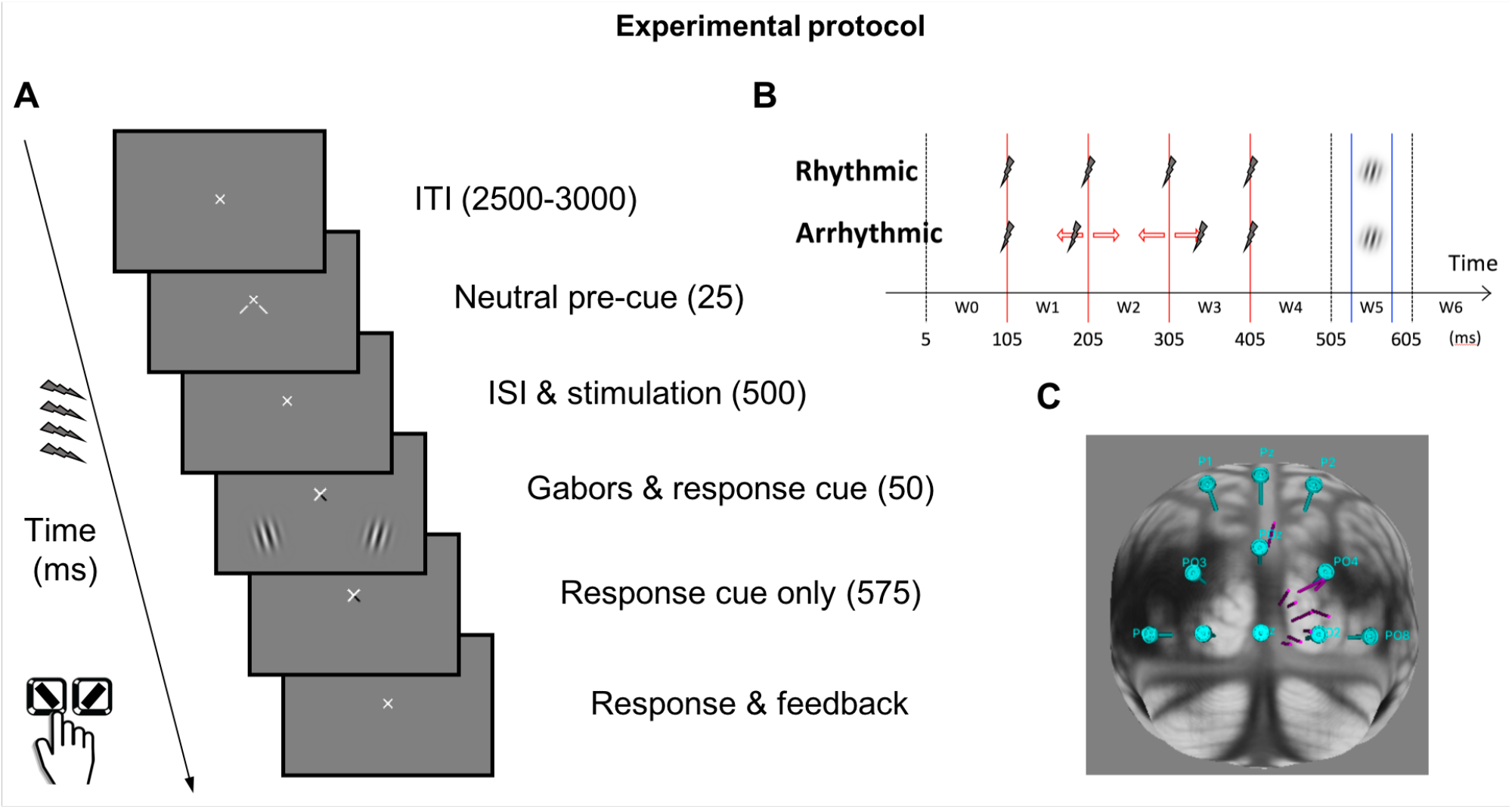
Experimental protocol (**A**), Trial structure. Each trial began with an intertrial period (ITI), followed by a neutral pre-cue to indicate the two stimulus locations. Active or sham stimulation was delivered in the interstimulus interval (ISI) between the neural pre-cue and the Gabor stimuli. The response cue indicated which Gabor was the target. Participants responded whether the target Gabor was tilted to the left or to the right with a key press. (**B**), Stimulation patterns & analysis time windows. The onset of the pre-cue is at t = 0. The four red lines mark the pulse timings in the rhythmic condition. The two blue lines mark the timings of Gabors onset and offset. In the rhythmic condition, the gap between pulses was exactly 100 ms. In the arrhythmic condition, the timing of the second and the third pulses was jittered (see text and Fig S1 for the probability distribution). Time window W0 is the cycle before the first pulse. Time windows W1, W2, and W3 are the cycles between pulse pairs. Time windows W4 and W5 are the first and the second cycles after the last pulse, respectively. (**C**), TMS loci & directions. Each cylinder represents the TMS directional vector of one participant on the MNI brain template. The cyan discs indicate the EEG electrode positions.

A central fixation cross was constantly present. Participants were asked to maintain fixation at all times and to blink after each trial. An eye tracker (EyeLink1000) ensured that fixation was within a 1.5° radius invisible circle. Trials for which fixation was broken (including blinking), from cue onset to stimuli offset, were discarded and repeated at the end of the block. The contrast level of the two Gabor patches were independently titrated before the experiment to attain a discrimination performance level of d’~1.2. A method of constant stimuli (4-80% Gabor contrasts in 7 log steps) was used to obtain the contrast at which sensitivity (d’)^71,72^ reached half-of-max sensitivity of the Naka-Rushton function was defined as c_50_ contrast. The group average (standard deviation) of c_50_ contrast was 16% (4%) for both visual fields.

During the ISI, 4 rhythmic or arrhythmic (**Fig. 1B**), active or sham pulses were delivered. The ISI between the pre-cue and the first pulse was 80 ms. The experiment applied a 2 (rhythmicity types) × 2 (stimulation types) within-subject design. The rhythmic and arrhythmic stimulations were blocked. The stimulation type (active or sham) and the target side were randomized within blocks. The experiment consisted of 8 blocks, each contained 64 trials, for a total of 512 trials; 128 trial repetitions per each of the 4 experimental conditions (2 rhythmicity types × 2 stimulation types). Should we find differences between the rhythmic-active and arrhythmic-active contrast but not between the rhythmic-sham and arrhythmic-sham contrast, this design would allow us to rule out TMS sound, rhythmicity, and differences between rhythmically and arrhythmically interpolated EEG signals as plausible explanations of the entrainment effect.

Given that during TMS stimulation the participants were engaged in a visual discrimination task, as a secondary analysis, we assessed whether visual performance was affected by entrainment. There were no significant effects regarding whether entrainment modulates visual discrimination^71,72^, either for visual sensitivity (d’; **Fig. S2**) or response criterion (c; **Fig. S3**). The error bars represent ±1 S.E.M. corrected for within-subject design^73^.

### Transcranial magnetic stimulation

The TMS site for each participant was defined by retinotopy or phosphenes (see Individualized TMS site). **Fig. 1C** illustrates the vectors connecting the cortical and scalp sites for each participant. All but one participant’s stimulation vector clustered around electrodes Oz, O2 and PO4. For that reason, we excluded that observer and replotted the data for the time frequency analysis. The TMS intensity was fixed at 70% for all participants (except for one at 65% and another at 67%) of the maximal machine output, to ensure no phosphene induction during the task. The fixed upper bound of the TMS intensity was independently determined in pilot studies to ensure that the coil would not overheat in the middle of an experimental block. The sham control consisted of 4 pulses of pre-recorded TMS sounds played through a speaker attached to the coil. The coil was placed over the scalp, targeting the same area regardless of the actual stimulation type. Participants were asked to report if they saw any phosphenes at any point during the task; they reported none.

In the rhythmic condition, the pulses were 100 ms apart, aiming to induce alpha-band entrainment. In the arrhythmic condition, the timing of the first and last pulses of the burst were the same as in the rhythmic condition, whereas the timing of the second and the third pulses were randomly jittered on each trial according to a bimodal distribution synthesized from normal distributions N (±30ms, 10ms) (**Fig. S1B**). Before and after the experiment, we verified that the registered TMS timings on the EEG achieved the expected precision (**Figs. S1C-D**).

### Individualized TMS site

We had magnetic resonance imaging (MRI) structural scan images and population receptive field (pRF) mapping^74,75^ data of the visual cortices for all but 3 participants. We used NiBabel^76^ to extract the right hemisphere V1 voxels corresponding to the location and size of the stimulus in the left visual field. The center of mass of the extracted voxels served as the individualized TMS target site. For the 3 participants without fMRI data, we used a TMS phosphene induction method^56,77–81^ to localize the optimal cortical site inducing unilateral phosphene perception in the left visual field (most likely on V1/V2^56,77,82–84^ and probably V3^85,86^). For these participants, the cortical site depth was arbitrarily set at 26.4mm, the average of the scalp-to-site distance of all other participants. **Table S1** summarizes the TMS cortical and scalp sites in Montreal Neurological Institute (MNI) coordinates. At these individualized stimulation sites defined by pRF voxels, all but 1 participant reported seeing phosphenes; thus, the pRF and the phosphene induction method yielded comparable stimulation sites.

### EEG preprocessing

The EEG recordings were digitized at 2500 Hz without any filtering. There was no re-referencing as the region of interest was the occipital cortex. The spike artifact caused by a TMS pulse typically lasted 4-10 ms. To reduce the artifact, signals within 1 ms before and 13 ms after each TMS pulse were replaced with shape-preserving piecewise cubic interpolated data (MATLAB command: interp1 with ‘pchip’ option)^43,48,57,87^. The exact same procedure was applied to EEG data during sham auditory pulses to ensure that any statistical differences could not be attributed to discrepancies in data processing. No trials were discarded other than those with blinks (see above, *Visual discrimination task*).

The continuous data were then down-sampled offline to 100 Hz and segmented into epochs containing data from 300 ms before to 900 ms after the neutral pre-cue onset. Preprocessing and analyses were carried out with MATLAB R2017a and software packages FieldTrip^88^ and Brainstorm^89^.

### EEG analyses

Before the experiment, we recorded 2-min eyes-closed resting-state activity to define the individual alpha frequency (IAF) as the frequency corresponding to the maximum peak between 7 and 13 Hz on Welch’s periodogram^90^ (MATLAB command: pwelch). To assess if the neural activity was phase-locked to the entraining periodic stimulation, we calculated two indices for each of the 4 conditions: (1) evoked oscillatory amplitude (square-root of power), by averaging waveforms across trials first and then applying Morlet wavelets^65^; (2) inter-trial phase coherence (ITPC, or phase-locking value)^91^ by applying Morlet wavelets to each trial and calculating their consistency with 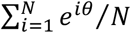, where i denotes the trial number, N the total number of trials, and θ the phase. ITPC is a ratio between 0 and 1, in which 1 indicates perfect phase alignment. For both indices, each Morlet wavelet had 5 cycles^57^, and the frequency range was 3-50 Hz. With 5 cycles of Morlet wavelets, the spectral bandwidth and wavelet duration, characterized by full width at half maximum, were 4.7 Hz and 0.2 sec, respectively. Thus, evoked oscillatory amplitude and ITPC centered around 10 Hz characterized the alpha-band (8-12 Hz) activity. Additionally, pre-TMS α-phase was defined as the phase calculated from Morlet wavelets, 200 ms before the first TMS pulse (95 ms before the pre-cue onset).

We analyzed the evoked oscillation amplitude and ITPC in 7 planned time windows: W0 (5-105ms, before the first pulse), W1 (105-205ms, between the first and second pulses), W2 (205-305ms, between the second and third pulses), W3 (305-405ms, between the third and last pulses), W4 (405-505ms, first cycle after the last pulse), W5 (505-605ms, second cycle after the last pulse), and W6 (605-705ms, third cycle after the last pulse). Because these time windows, which are consistent with reported time intervals of entrainment effects by TMS^43,48,57^, were specified a priori, multiple comparison correction was not required. But within each time window topographic analyses (paired t-tests between conditions across all channels) were corrected for multiple comparison with cluster-based permutation tests^92^ (1000 permutations, one-tailed, alpha=.05, cluster alpha=.05). According to the trial structure of the visual discrimination task, stimuli were presented in the middle of W5 (530-580 ms). Online eye tracking ensured that the trials containing eye blinking were discarded so that the analyzed EEG time windows were free of such artifacts.

Several statistical tests were performed. (1) As entrainment should lead to localized elevation of evoked oscillation amplitude and ITPC near the stimulation site, t-tests were performed by averaging the neural signatures in channels O2 and PO4 in the 10-Hz band at all time windows. (2) To explore the temporal and spectral specificity of entrainment, t-tests of the two neural signatures were performed across all time-frequency bins averaged across channels O2 and PO4. (3) To explore the topography of neural activity before, during and after entrainment, t-tests were performed across all channels in the 10 Hz band at all time windows. (4) To examine whether ITPC enhancement following rhythmic TMS depended on pre-TMS alpha phase, trial-by-trial 10-Hz phases 200 ms before the first pulse (95 ms before the pre-cue onset) were assorted into 6 equidistant phase bins. ITPC across participants and time points in W2-W6 indexed by these 6 phase bins were analyzed via one-way repeated measures ANOVA and a regression analysis with a sine wave (y = a*sin(f*x*π/3+ϕ)+c, where x is the bin number; a, f, ϕ, and c were free parameters)^57^. (5) To examine the relation between IAF and ITPC, a linear regression was performed.

For the rhythmic-active condition, the arrhythmic-active condition is a more stringent control than the rhythmic-sham condition across all figures (**Figs. 2–5, S6-S9**) (see also ^57^). Across all time windows (**Figs. 3–5, S6-S9**), the contrasts between the rhythmic-active and arrhythmic-active conditions were significant whenever those between the rhythmic-active and rhythmic-sham condition were significant, except for evoked oscillation amplitude and ITPC in time window W0 (the 100 ms cycle before the first pulse) and evoked oscillation amplitude in time window W1 (the 100 ms cycle after the first pulse) (**Figs. S6, S7A**). Therefore, we report the statistical contrast between the rhythmic and arrhythmic-active stimulation conditions.

**Figure 2.**
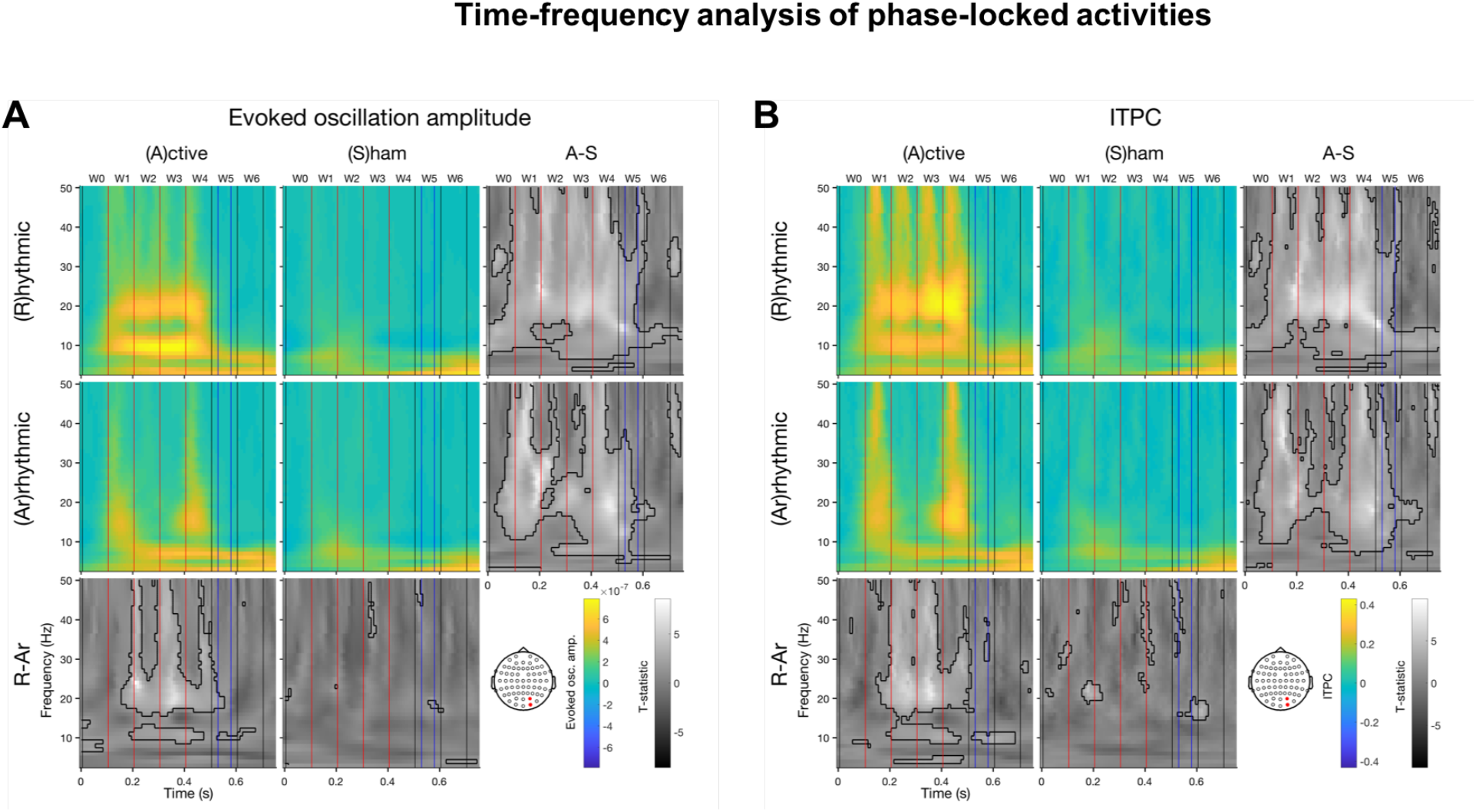
Time-frequency analysis of phase-locked activities. Colored panels are the group-averaged activities per condition (rhythmic vs. arrhythmic × active vs. sham stimulation). Gray panels are the t-statistics of condition contrasts per column or row. The pre-cue onset was defined as t=0. The electrode of interest is the average of the two channels (O2 & PO4) near the TMS loci. In the t-statistic panels, the thick black contours indicate significant time-frequency bins. In each panel, the vertical black and red lines demarcate the 100 ms time windows, blue lines the Gabor onset and offset. The red lines also indicate the pulse timings in the rhythmic conditions. (**A**), evoked oscillation amplitude. (**B**), ITPC. Each TMS pulse elicits broad band responses. Rhythmic-active stimulation elicits activities around 10 Hz and the first harmonic. Note that the arrhythmic-active condition is a more stringent control than the rhythmic-sham condition for the rhythmic-active condition. Note that at 10 Hz, the full width at half maximum of time and frequency for Morlet wavelets with c=5 are 0.19 and 4.71, respectively.

**Figure 3.**
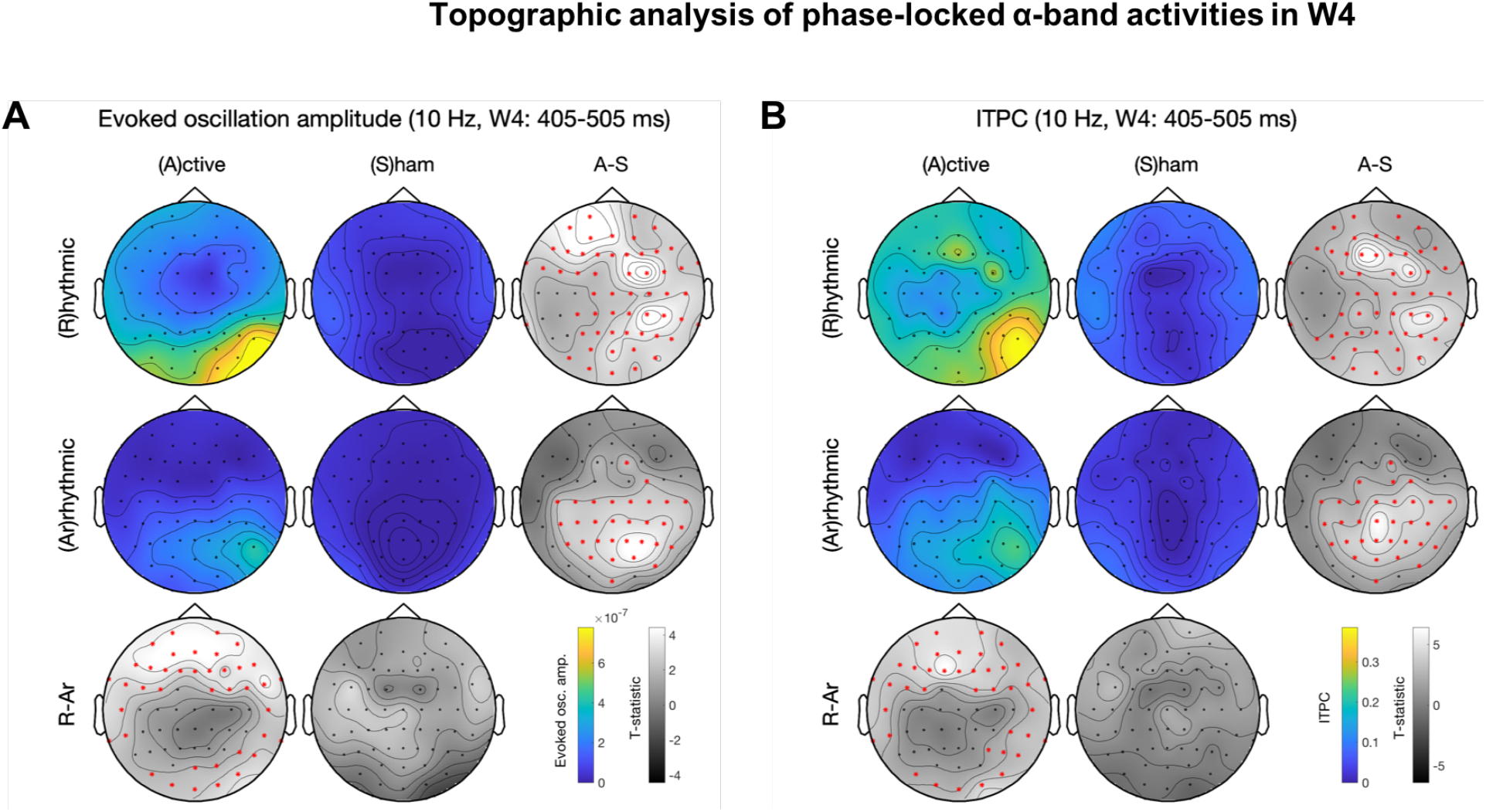
Topographic analysis of phase-locked α-band activities in W4. The panel layout is similar to that in **Fig. 2**. The red star symbols indicate significant channels after cluster-based permutation test for multiple comparison correction. (**A**), evoked oscillation amplitude. (**B**), ITPC. Rhythmic-active stimulation elicits widespread activation. With arrhythmic-active stimulation as control, there are significant clusters in the frontal and the occipital regions. Note that the patterns between (**A**) and (**B**) are similar.

## Results

We present psychophysics results, ERP analyses, wavelet analyses relevant for each of the 4 hypotheses, followed by exploratory analyses regarding the temporal dynamics of the topography of the entrainment effect. Altogether both sets of results indicate successful entrainment.

### Psychophysics

Visual sensitivity (d’) and criterion (c) in a discrimination task was unaffected by the elevated alpha-band evoked oscillation amplitude and ITPC (**Figs. S2** and **S3**).

### ERP

The spline interpolation method reasonably cleaned up the TMS spike artifacts present in the ERPs (**Fig. S4**). Apart from wavelet analyses, we carried out alpha-band bandpass filtering of ERPs across conditions (**Fig. S5**) and noted: 1) intrinsic alpha oscillations are evident before either active or sham stimulation, 2) in the active conditions, the pulse timings are closely aligned with alpha wave peaks, and 3) the intervals between waveform peaks are close to 100 ms only in the rhythmic-active condition.

#### (1) Progressive enhancement of entrainment

To evaluate the temporal evolution of entrainment, we obtained t-test results of the planned contrasts between rhythmic and arrhythmic-active stimulation conditions in the 7 pre-defined time windows (**Materials and Methods**) for evoked oscillation amplitude and ITPC (**Table 1**). Consistent with our prediction regarding entrained neural activity, the planned contrasts were not statistically significant 100-ms before and after the first TMS pulse (W0 and W1, p>.05), but became significantly different for the later time windows (W2, W3, and W4, p<.05). Interestingly, the occipital alpha-band stimulation enhanced phase-locked activity during the second and the third cycles after the last pulse (significant contrasts in W5 and W6, p<.05). Our entrainment effect in terms of both evoked oscillation amplitude and ITPC lasted for 2-3 cycles (200 and 300 ms, respectively, after the last pulse of the TMS burst).

**Table 1.**
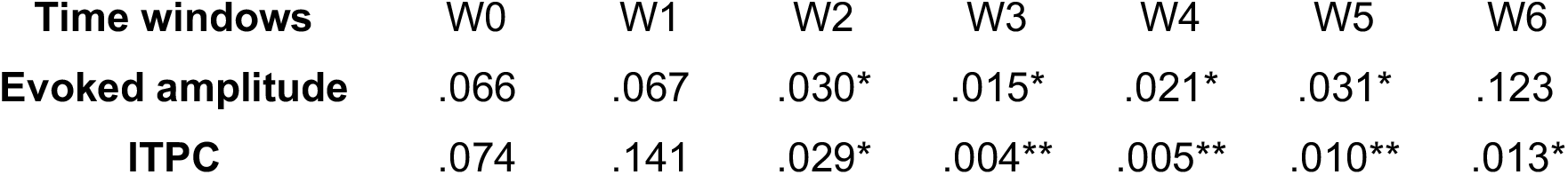
p values of planned t-tests of evoked oscillation amplitude and ITPC at the α-band. The frequency of interest is 10 Hz. The electrode of interest is the average of O2 and PO4 (near the stimulation loci; see **Fig. 1C**). The planned contrasts are between rhythmic and arrhythmic-active stimulation conditions in the 7 pre-defined time windows, including the periods before (W0), during (W1-W3), and after (W4-W6) stimulation. The contrasts are significant starting from W2 until W5 or W6. *: p<.05, **: p<.01

| Time windows | W0 | W1 | W2 | W3 | W4 | W5 | W6 |
| --- | --- | --- | --- | --- | --- | --- | --- |
| Evoked amplitude | .066 | .067 | .030* | .015* | .021* | .031* | .123 |
| ITPC | .074 | .141 | .029* | .004** | .005** | .010** | .013* |

#### (2) Output frequency specificity of entrainment

To assess the output frequency-specificity, we conducted time-frequency analysis of phase-locked activities with frequencies ranging 3-50 Hz and time ranging 0-750 ms (**Fig. 2**). Rhythmic-active stimulation elicited activity at ~10 Hz and its first harmonic (~20 Hz). Additionally, there were broad band responses for each of the 4 TMS pulses. Contrasting the rhythmic-active condition with either the arrhythmic-active or the rhythmic-sham condition revealed that only the ~10 Hz rather than the first harmonic component lasted beyond 1 cycle after the last TMS pulse, demonstrating output frequency specificity. Overall, the patterns for evoked oscillation amplitude and ITPC (**Fig. 2A,B**) were similar.

#### (3) Dependence on the intrinsic oscillation phase

To ensure that the enhanced phase-locked activity reflected entrainment rather than reverberation of an externally imposed rhythm, we performed regression analyses on ITPC across the time windows that were significant (W2-W6) in **Table 1**. The pre-TMS phases (200 ms before the first 10-Hz pulse) were assorted into 6 equidistant phase bins^57^. One-way repeated measures ANOVA revealed that the ITPC values across phase bins were significantly different [*F*(9,45)=6.67, *p*<1e-5]. In this period, ITPC depended on the phase bin with a sinusoidal trend (**Fig. 6A)**, as previously reported^57^, suggesting that the phase of ongoing alpha oscillations matters. This result refutes the reverberation account, according to which ITPC would have been a flat line.

#### (4) Input frequency dependence of entrainment

Inconsistent with the hypothesis that the intrinsic oscillators operate at the IAF in the occipital cortex, the degree of similarity between IAF and 10 Hz did not correlate with entrained ITPC [*r*=.04, *p*=.91] (**Fig. 6B**). Instead, IAF directly correlated with entrained ITPC [*r*=.63, *p*<.05] (**Fig. 6C**). The degree of similarity between IAF and 10 Hz did not correlate with evoked oscillation amplitude [*r*=.15, *p*=.68] (**Fig. S10A**). IAF did not correlate with evoked oscillation amplitude [*r*=.39, *p*=.27] (**Fig. S10B**). Moreover, pre-TMS total alpha oscillation amplitude did not predict entrained ITPC (**Fig. S15**).

### Further analyses

Similar pattern of results were obtained for each of the previous analyses, when (a) the observer whose stimulation location (near POz) differed from the rest was excluded (**Figs. S11-S12 and Table S2**), and (b) taking only the trials that were preceded by correct responses (76% of the data) (**Figs. S13-S14 and Table S3**). These results show that the effect generalizes across observers and that the feedback for incorrect responses did not drive the differential responses among the 4 conditions (rhythmic-sham, arrhythmic-sham, rhythmic-active and arrhythmic-active conditions).

### Temporal dynamics of 10-Hz topography

We explored the topography of the entrainment effects by performing topographic analysis of phase-locked alpha-band activities in each time window and assessing the temporal dynamics of evoked oscillation amplitude and ITPC. Cluster-based permutation tests^92^ were used for correction of multiple comparisons. Descriptively, with the same alpha threshold for cluster-based permutation test, 10-Hz activity was initially widespread, likely due to EEG volume conduction, and progressively became more local over time. The statistically significant cluster included the frontal and occipital regions for both evoked oscillation amplitude and ITPC in W2-W4 (**Figs. 3, S8-S9**). Note that the occipital clusters in W2-W6 (**Figs. 3–5, S8-S9**) were more lateralized towards the stimulated (right) side.

A genuine entrainment effect should start in later time windows because this phenomenon has been characterized as a gradual process that realigns the intrinsic oscillator’s phase (hence phase dependent, **Fig. 6A**)^57^. The hypothesis of progressive enhancement of entrainment was corroborated by the fact that the topographic contrast between rhythmic-active and arrhythmic-active stimulation was not significantly different for either evoked oscillation amplitude or ITPC in W0 (**Fig. S6**) and for evoked oscillation amplitude in W1 (**Fig. S7**). ITPC was significantly different during W1-W4 (**Figs. 3, S7-S9**) and continued to be so after evoked oscillation amplitude was no longer different in W5-W6 (**Figs. 4–5**). This pattern of results based on topography paralleled the results from planned t-tests nearby the stimulation site (**Table 1**).

**Figure 4.**
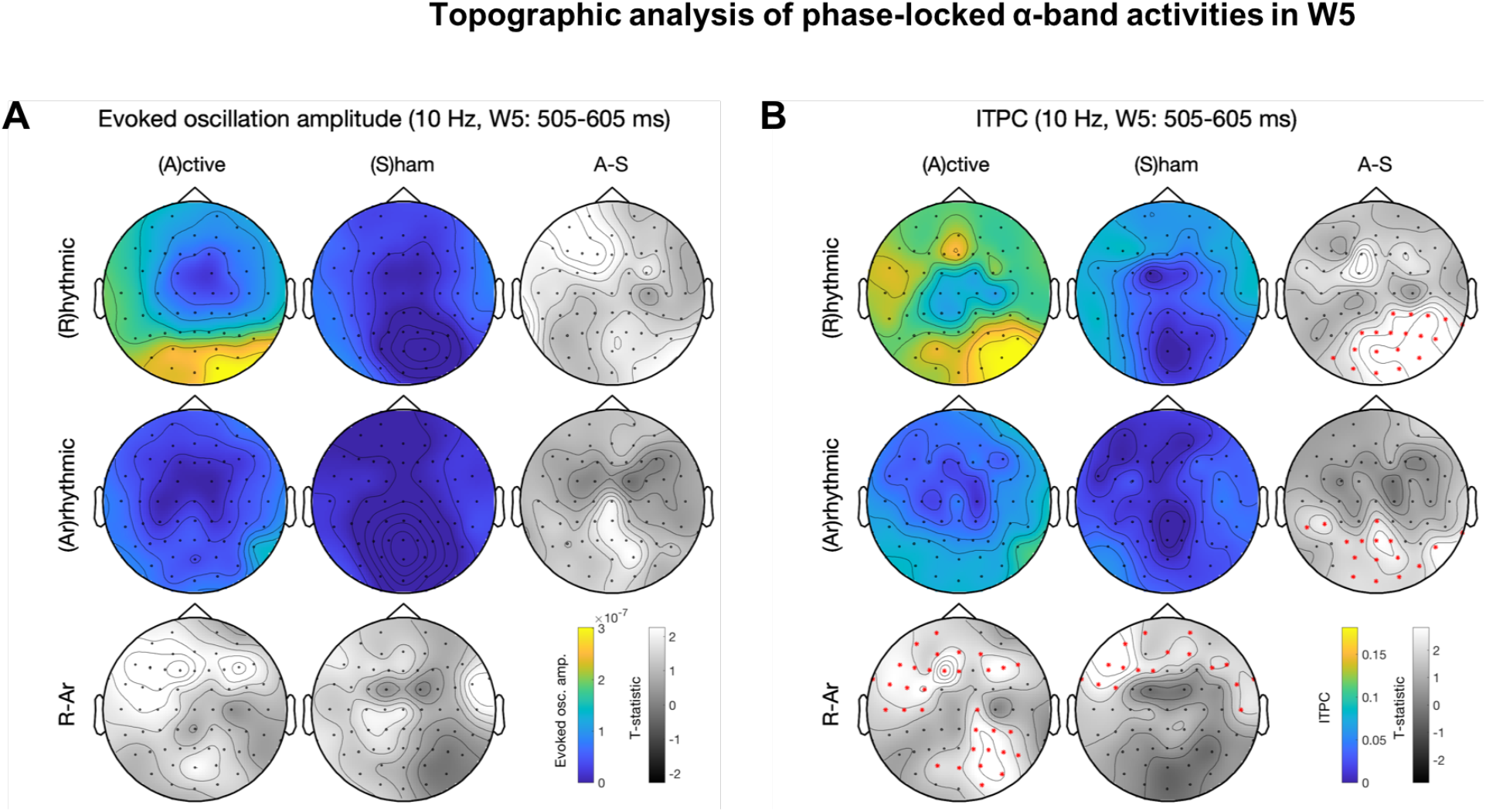
Topographic analysis of phase-locked α-band activities in W5. See **Fig. 4** for panel layout description. (**A**), evoked oscillation amplitude. (**B**), ITPC. Rhythmic-active stimulation elicits widespread activation. Note that (**B**) continues to show significant difference between conditions whereas (**A**) does not.

## Discussion

This study is the first to show TMS entrainment of alpha activity in occipital cortex. We evaluated whether occipital alpha-band activity can be entrained by TMS targeting V1/V2 while participants performed a visual discrimination task. We tested the following hypotheses, based on established entrainment characteristics: (1) *progressive enhancement of entrainment*, (2) *output frequency specificity*, (3) *dependence on the intrinsic oscillation phase*, and (4) *input frequency specificity*. Our results are consistent with the first three hypotheses, revealing that occipital alpha activity can be entrained. However, the results were not consistent with the 4^th^ hypothesis. Instead, IAF correlated with ITPC.

### Successful and lasting alpha-band entrainment in the occipital cortex

With a 4-pulse 10-Hz TMS protocol, we successfully entrained the intrinsic alpha oscillations in the right occipital cortex for 3 cycles (300 ms) after the last TMS pulse. This novel result further supports that short-burst rhythmic TMS can effectively entrain frequency-specific neural synchrony, trial-by-trial, in the stimulated region. To our knowledge, this is the most effective short-burst TMS entrainment finding up to date. Previous TMS-EEG studies using a similar short-burst (3-5 pulses) rhythmic stimulation and control conditions achieved entrainment of intrinsic oscillations for about 1-1.5 cycles after the last pulse, with or without a concurrent visual task (**Table 2**). We may have obtained longer lasting TMS-entrained duration than the study entraining right IPS^57^ because our participants were engaged in a visual task, instead of in resting state. The only study evaluating TMS entrainment in occipital cortex reported not to find it^48^, even though their participants were also engaged in a visual discrimination task; their 3 10-Hz TMS pulses may have been insufficient for entraining the occipital cortex. Future studies may further investigate which factors are crucial to successfully induce entrained synchronization of intrinsic neural oscillations outlasting rhythmic stimulation.

**Table 2.**
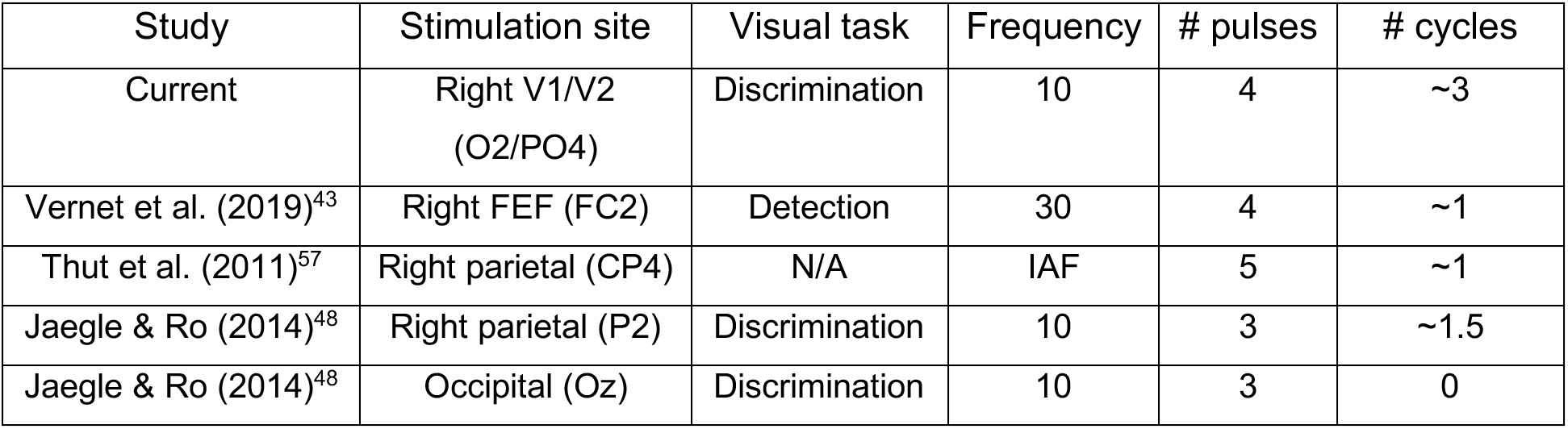
Comparison with previous short-burst TMS-EEG entrainment studies. The stimulation site lists the target brain region along with the closet EEG electrode. The visual task is the task concurrent to TMS and control conditions. Most studies focus on the intrinsic alpha oscillations in the occipital or parietal cortices except one on the intrinsic beta oscillations in the frontal cortex. The number of cycles indicates how long the entrainment effect lasts following the last TMS pulse.

Note that our results fulfilled entrainment requirements and ruled out a reverberation account (as in ^57^). Moreover, the evoked oscillation amplitude and ITPC at the alpha-band were not significantly different between the rhythmic and arrhythmic-active stimulation conditions in time windows W0 and W1. Therefore, it is unlikely that the lasting effects we found were due to temporal leakage of wavelets. The number of wavelet cycles (c=5) in our study was the same as in the first concurrent TMS-EEG entrainment study^57^, where only ~1 cycle of alpha-band entrainment outlasting the last stimulation pulse was reported in posterior parietal region. Our design allowed us to rule out TMS sound, rhythmicity, and differences between rhythmically and arrhythmically interpolated EEG signals as plausible explanations of the entrainment effect.

Our results reveal that changes of two phase-locked activity measures –evoked oscillation amplitude and ITPC– show different temporal dynamics beyond the stimulation burst. ITPC lasted one more cycle than evoked oscillation amplitude (**Table 1**; **Figs. 2,4,5**). As ITPC is a phase-locking activity measure that does not take amplitude into account, this finding suggests that phase could be more sensitive and informative than amplitude to index the occurrence and duration of entrainment effects in the occipital cortex.

**Figure 5.**
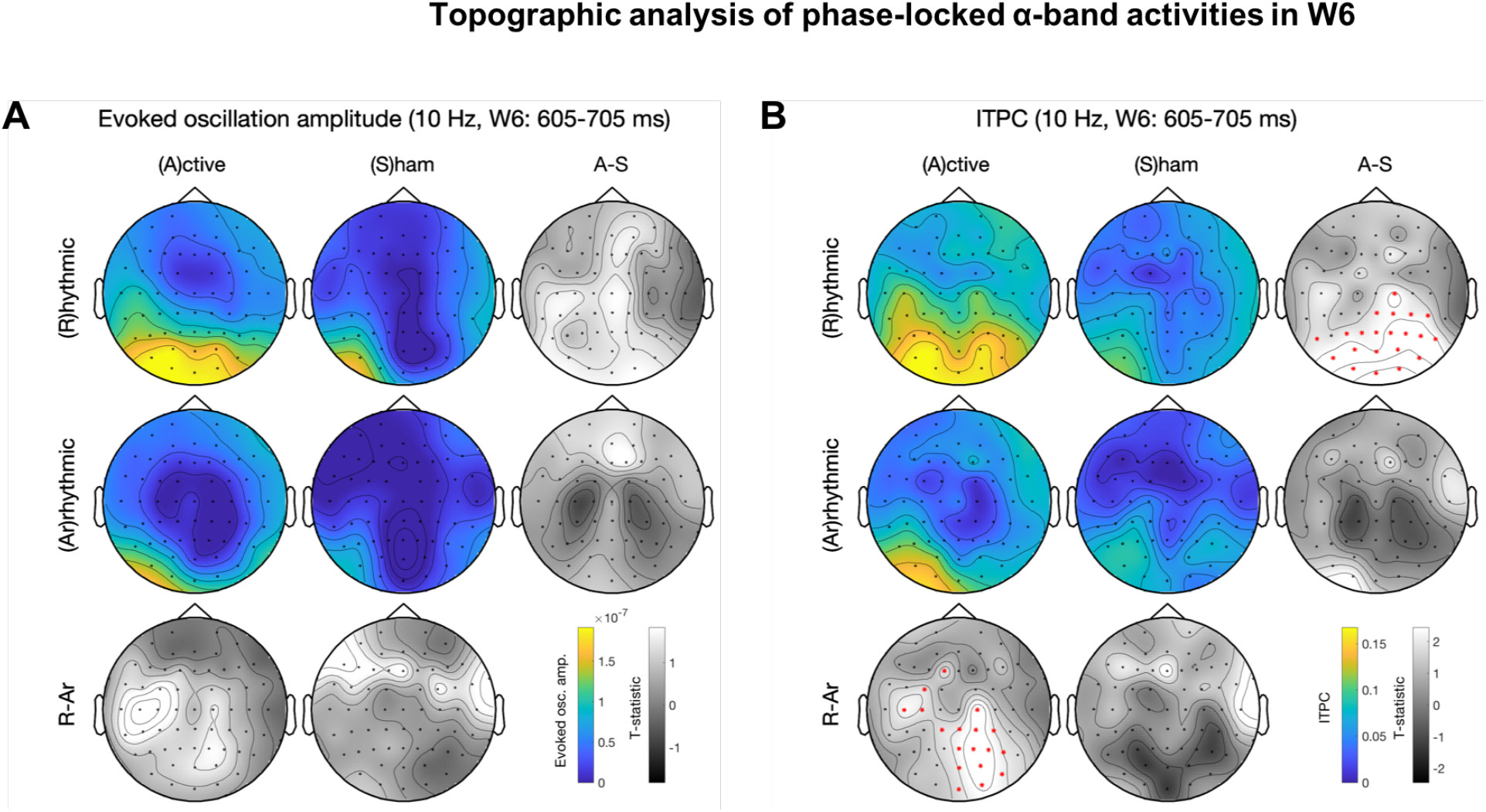
Topographic analysis of phase-locked α-band activities in W6. See **Fig. 4** for panel layout description. (**A**), evoked oscillation amplitude. (**B**), ITPC. Rhythmic-active stimulation elicits widespread activation. Note that (**B**) continues to show significant difference between conditions whereas (**A**) does not. A significant lateralized parieto-occipital cluster exists in the rhythmic- vs. arrhythmic-active stimulation contrast.

**Figure 6.**
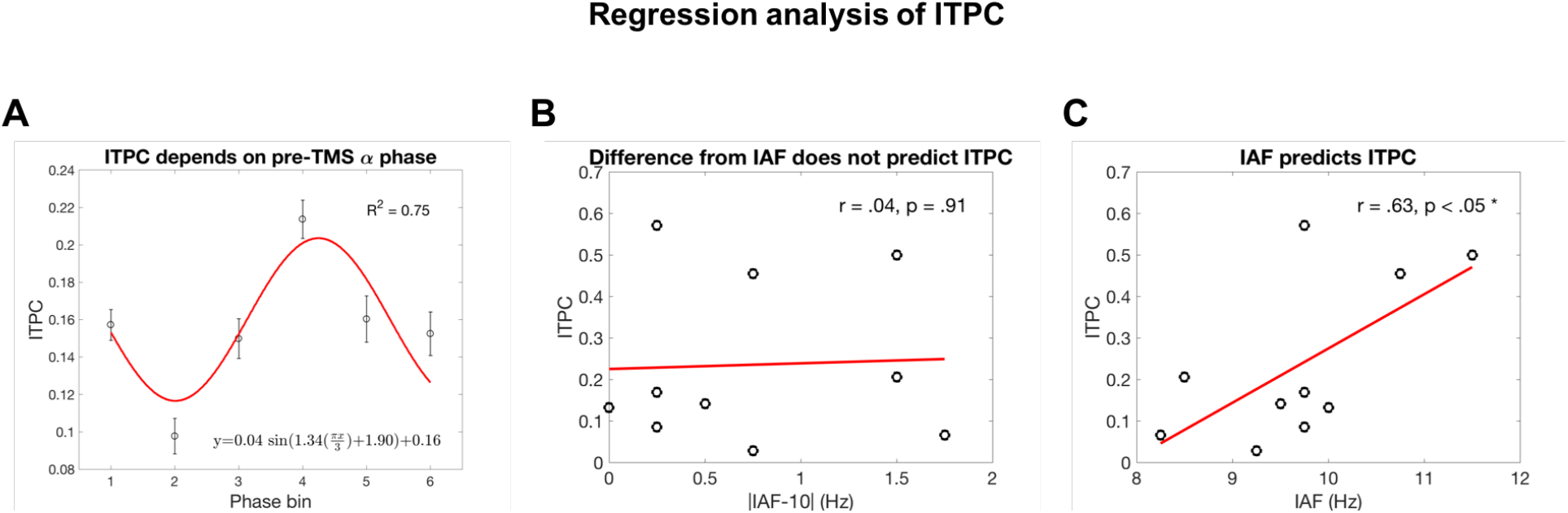
Regression analysis of ITPC in the rhythmic-active stimulation condition. The time window of ITPC is 0.205-0.705 s, the same as the significant windows in **Table 1**. (**A**), ITPC depends on pre-TMS α-phase (200 ms before the first pulse). Each point is the average of 510 samples (10 participants × 51 time points). The error bars represent ±1 SEM. This ~1 period cyclic pattern suggests that the congruency between TMS and the phase of ongoing α oscillations determines the effectiveness of entrainment. Phase bin #2 roughly corresponds to the ~10 Hz ERP peak and phase bin #5 to the trough. Note that due to conduction time the phase measured on the scalp does not indicate the exact phase of the oscillator in the brain. (**B**), Difference from IAF does not predict ITPC. The absolute difference between IAF and the entraining frequency does not predict ITPC during and after stimulation. (**C**), IAF predicts ITPC. IAF positively correlates with ITPC during and after stimulation. See text for discussion.

The finding that the occipital cortex can be entrained by TMS is consistent with the notion of inter-areal entrainment as a form of neural communication whose function could be to achieve local phase alignment^36,49^. Given that visual stimuli were presented at the same timing across all conditions, the continued phase-locking in the second and third cycles after the last pulse (W5 and W6) cannot be attributed to evoked alpha activity by target onset.

We note that during 10 Hz TMS (W1-W4) there were clear first harmonic (20 Hz) responses both in terms of evoked oscillation amplitude and ITPC (**Fig. 2**). However, after TMS (W5-W6) only the 10 Hz evoked oscillation amplitude and ITPC were long lasting. These lasting responses support the notion of entrainment. Furthermore, although our auditory sham did not produce a tactile stimulation, we note that this difference cannot explain the entrainment in the occipital cortices because a tactile entrainment would have been contralateral to TMS.

### IAF predicts occipital entrainment

Some studies have assumed that using IAF, the intrinsic oscillation frequency of the parieto-occipital cortex at rest, as the entraining frequency would be optimal for entrainment, but this assumption has rarely been tested^37^. Similarly, our TMS-EEG findings are inconsistent with such an assumption: The dependency of ITPC on pre-TMS phase shows that entrainment effects depend on the ongoing oscillatory state while participants are engaged in a visual task, refuting a passive reverberation account (**Fig. 6A**). Therefore, intrinsic alpha oscillators exist in the occipital cortex. Furthermore, pre-TMS alpha power could not reliably predict entrained ITPC (**Fig. S15**). These findings are consistent with those of a transcranial alternating current stimulation (tACS)-EEG study^93^ showing that neither the baseline power nor IAF could reliably predict alpha power increase after 10-Hz tACS entrainment.

The reason that the difference between the entraining frequency and IAF did not modulate ITPC (**Fig. 6B**) is likely the high stimulation intensity of TMS. Consistent with the idea of Arnold tongue (synchronization region defined on an amplitude-to-frequency plane) for oscillatory systems^94^, it has been shown that when the amplitude of the entraining stimulation is above a certain threshold, the output oscillating frequency is at a frequency equal to the stimulation instead of the intrinsic frequency^95^. With increasing intensity, a wider range of stimulation frequency centered around the intrinsic frequency could result in reliable entrainment as evidenced by the magnitude of phase-locking^96^.

The positive correlation between IAF and ITPC (**Fig. 6C**) suggests that individuals with higher IAF may be more susceptible to occipital alpha-band entrainment. A possible explanation is that the higher the IAF, the shorter the oscillation period. In response to external stimulation, shorter periods imply smaller absolute time steps in phase re-alignment. Future experimental studies may further exploit this interesting relation and examine if the same conclusion holds with IAF estimated during eyes-open resting state. This finding may also provide constraints for future cortical network modeling studies.

It is an open question whether the finding that IAF in the occipital cortex correlates with locally TMS-entrained synchrony can be generalized to other brain areas, such as the parietal cortex or the somatosensory cortex. Interestingly, the closer the stimulation frequency to the individual beta frequency in the primary motor cortex, the more entrained the phase-locking^46^. Together with our findings, these results suggest that alpha and beta oscillations subserve distinct functions or that sensory and motor cortices have different operating principles.

### Can alpha-band occipital TMS entrainment modulate visual sensitivity (d’)?

Our results show that visual sensitivity and criterion in a discrimination task was unaffected by the elevated alpha-band evoked oscillation amplitude and ITPC (**Figs. S2** and **S3**). Given that we only tested one specific target presentation timing, and at one TMS intensity per participant, our result does not necessarily rule out the possibility that changes in phase-locked activity level could modulate d’. For instance, a study showed that hit rate, but not false alarm rate, in the visual field ipsilateral to TMS was significantly higher immediately after a 5-pulse alpha-band TMS over the occipital cortex than that in the sham condition^38^. Likewise, after alpha-band visual flicker, discrimination accuracy oscillates ~10 Hz^97,98^. Thus, the testing of multiple time lags at multiple intensities, as well as entraining other locations (e.g., electrode CP4), stimulating at different frequencies (e.g., theta), or manipulating spatial attention^55,56,77^ could inform whether accuracy oscillates after occipital TMS entrainment and whether more elevated evoked oscillation amplitude and ITPC are required for a significant behavioral outcome.

In contrast to phase-locked analyses, some visual detection studies with trial-by-trial spontaneous oscillation analysis have revealed that pre-stimulus alpha power^99,100^ and phase^101^ correlate with response criterion (c) instead of visual sensitivity (d’). Further, for visual discrimination tasks, pre-stimulus alpha power correlates with confidence instead of accuracy^102^. However, note that without entrainment, trial-by-trial spontaneous pre-stimulus alpha activity is random and hence non-phase-locked across trials. Therefore, the current findings do not necessarily negate the possibility that entrainment may alter d’ following alpha-band entrainment.

### Protocol considerations for future studies

An advantage of our protocol is that it has an arrhythmic-active stimulation control condition, similar to studies investigating IPS^57^ and FEF^43^ areas. Across time windows, using the arrhythmic-active and rhythmic-sham control conditions led to similar statistical outcomes; overall, the arrhythmic-active condition was a more stringent control condition resulting in less widespread significant statistical differences at the scalp level (**Figs. 2–5, S6-S9**). Therefore, follow-up studies may consider removing the sham conditions to increase the statistical power of different types of TMS trials.

To further strengthen the importance of entraining near the intrinsic oscillation frequency, future studies could evaluate whether distinct TMS frequencies produce long lasting effects in other frequency bands. For instance, given our present findings establishing entrainment in the occipital cortex with TMS, the same protocol could be tailored to examine the theta band (4-7 Hz), which has also been reported during attention tasks^77,103^.

Our 4-pulse alpha-band TMS protocol provides an interesting alternative for occipital entrainment to that of tACS protocols. Alpha-band tACS protocols typically involves 10-20 minutes of stimulation followed by 1-3 minutes of testing period, during which enhanced alpha-power or ITPC has been recorded with concurrent EEG and taken as evidence of entrainment^93,104^ (but see alternative interpretations^105,106^). In any case, TMS protocols^48,57^, including ours, provide more focal stimulation effects than tACS and can be effectively delivered in shorter ‘bursts’ of pulses, hence pinpoint more accurately the temporal dynamics of entrainment effects, enable trial-by-trial stimulation designs and concurrent EEG recordings.

Entrainment helps synchronization across neural populations and communication in the brain, and supports cognitive processes such as top-down attention^40,42,43,49–52^. External manipulation protocols via TMS can help establish the causal role of intrinsic oscillations within a cortical area on specific brain processes and behaviors, and their interactions with other brain regions. Regarding translational implications, the established causality could be conducive to identify abnormal rhythms and for potential interventions. Specific constraints discovered with one specific brain stimulation technology, such as TMS, may be also informative to other technologies, such as tACS.

To conclude, we have established a rhythmic 4-pulse alpha-band occipital TMS protocol for effective trial-by-trial, online brain stimulation to enable alpha entrainment in retinotopically organized V1/V2 areas. With a 300-ms entraining period, phase-locked activities persisted for ~300 ms (three 10-Hz cycles) after the last pulse. To our knowledge, this is the longest lasting short-burst TMS entrainment finding reported up to date. Moreover, we found that IAF predicts the strength of entrained phase-locking across trials (ITPC). Therefore, IAF is a key factor worth investigating in future alpha entrainment studies.

## Acknowledgements

This research was supported by National Institute of Health (NIH R21-EY026185 and R01-EY019693) to M.C., IHU-A-ICM-Translationnel, ANR projet Générique OSCILOSCOPUS and Flag-era-JTC-2017 CAUSALTOMICS to A.V.-C. We thank Antonio Fernández and Zhilin Zhang for assistance in data collection; Noah Benson and Marc Himmelberg for guidance on fMRI retinotopy data analysis; Antonio Fernández and Chloé Stengel for valuable discussions regarding this project; Rachel Denison and Florencia Assaneo for useful comments on the manuscript.

## Author Contributions

Y.-J.L., L.D., A.V.-C., and M.C. designed the study. Y.-J.L. and L.S. developed software and performed the experiment. Y.-J.L. analyzed data and prepared figures. Y.-J.L. wrote the manuscript (with guidance and supervision from M.C.). L.D., A.V.-C., and M.C. edited the manuscript. A.V.-C. and M.C. supervised the project. M.C. administered the project and acquired the funding.

