## Supplementary materials for "Transcranial magnetic stimulation entrains alpha oscillatory activity in occipital cortex"

**Table S1. Stimulation loci.** The stimulation loci are individualized by fMRI pRF mapping, phosphene induction, or both. The # symbols indicate that the pRF mapping was unavailable and the TMS loci were determined by phosphene induction instead. The phosphenes were short lasting spatially circumscribed percepts reported in the hemifield contralateral (left) to the stimulated occipital pole (right).

| Participant | Scalp MNI coordinate | Target MNI coordinate | Distance to skin (mm) | TMS error (mm) |
|---|---|---|---|---|
| 01 | 23.33, -109.39, 1.75 | 16.44, -84.57, -4.84 | 28.6 | N/A |
| 02 | 13.62, -111.94, -1.65 | 10.06, -93.29, -3.08 | 18.8 | 0.25 |
| 03 | 13.75, -110.00, -2.74 | 4.75, -81.60, -4.68 | 31.4 | 0.32 |
| 04 | 6.96, -112.80, 2.17 | -1.25, -72.34, -7.93 | 42.5 | 0.52 |
| 05 | 11.84, -109.25, 5.02 | 6.41, -81.99, 4.22 | 28.9 | 0.48 |
| 06 | 25.50, -109.04, 6.42 | 19.48, -92.22, 4.63 | 18.1 | N/A |
| 07 | 1.28, -96.55, 37.55 | 0.30, -82.20, 29.43 | 16.5 | 0.52 |
| 08# | 5.23, -90.72, 2.81 | 11.21, -115.17, 10.11 | (26.4) | 0.58 |
| 09# | 12.95, -80.80, 3.59 | 25.17, -102.48, 11.45 | (26.4) | 0.76 |
| 10# | 5.87, -88.22, 7.32 | 17.92, -111.63, 11.68 | (26.4) | 0.46 |

**Table S2. p values of planned t-tests of evoked oscillation amplitude and ITPC at the α-band.** In this analysis, the participant whose stimulation site (near POz) deviated from the sites of others was excluded (n=9). The statistical patterns are the same as in **Table 1** (except for evoked oscillation amplitude in W5, which is now marginal). *: p<.05, **: p<.01

| **Time windows** | W0 | W1 | W2 | W3 | W4 | W5 | W6 |
|---|---|---|---|---|---|---|---|
| **Evoked amplitude** | .096 | .084 | .045* | .029* | .041* | .063 | .246 |
| **ITPC** | .068 | .078 | .027* | .008** | .001** | .018* | .025* |

**Table S3. p values of planned t-tests of evoked oscillation amplitude and ITPC at the α-band.** In this analysis, trials following incorrect responses were removed (~24% of total trials). The patterns are the same as those in **Table 1**, indicating that carry-over effects from the previous trial play little role in the entrained effects. *: p<.05, **: p<.01

| **Time windows** | W0 | W1 | W2 | W3 | W4 | W5 | W6 |
|---|---|---|---|---|---|---|---|
| **Evoked amplitude** | .356 | .142 | .039* | .011* | .010** | .017* | .147 |
| **ITPC** | .481 | .287 | .038* | .006** | .003** | .002** | .005* |

### Precision of stimulation timing

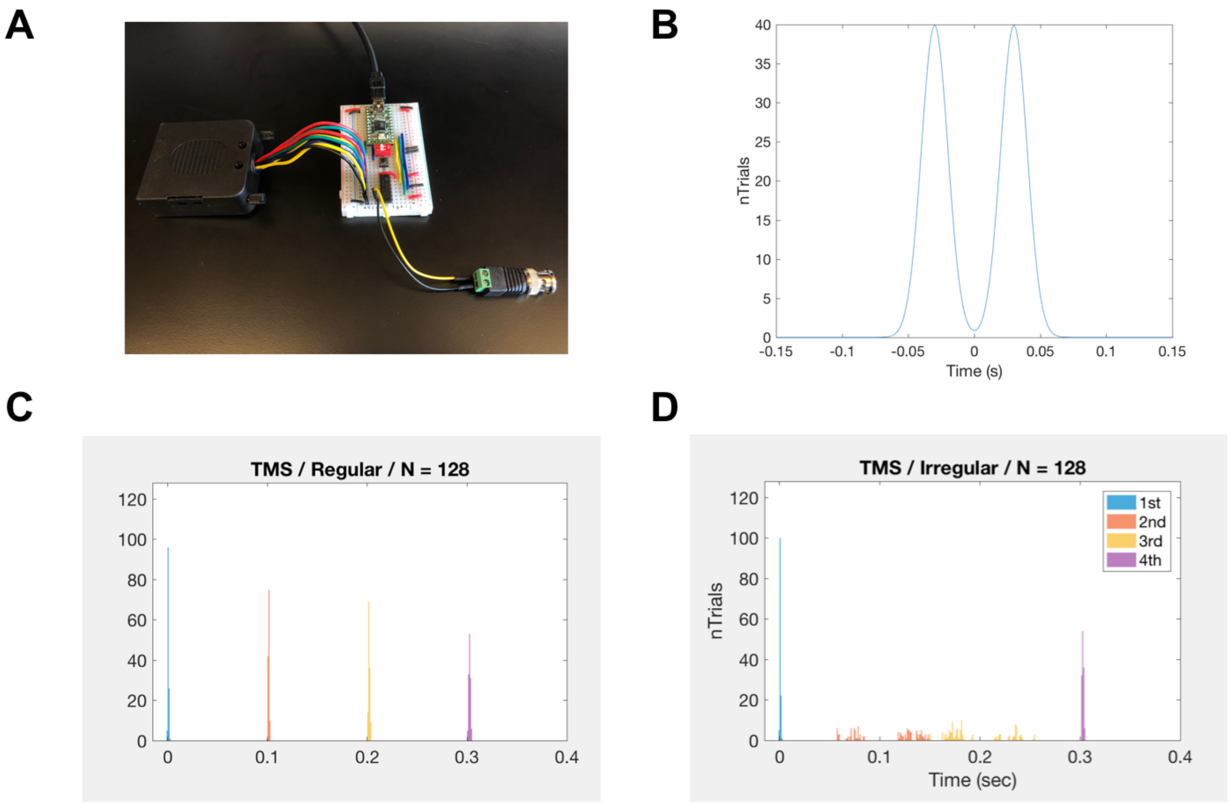


**Figure S1.** Precision of stimulation timing. (**A**), The MarkStim TMS triggering and EEG event registering device. (**B**), The timing jittering function. The second and the third pulses of the arrhythmic condition follows this function for temporal jittering. (**C**), Verified TMS timings in the rhythmic and (**D**), in the arrhythmic conditions of a representative participant.

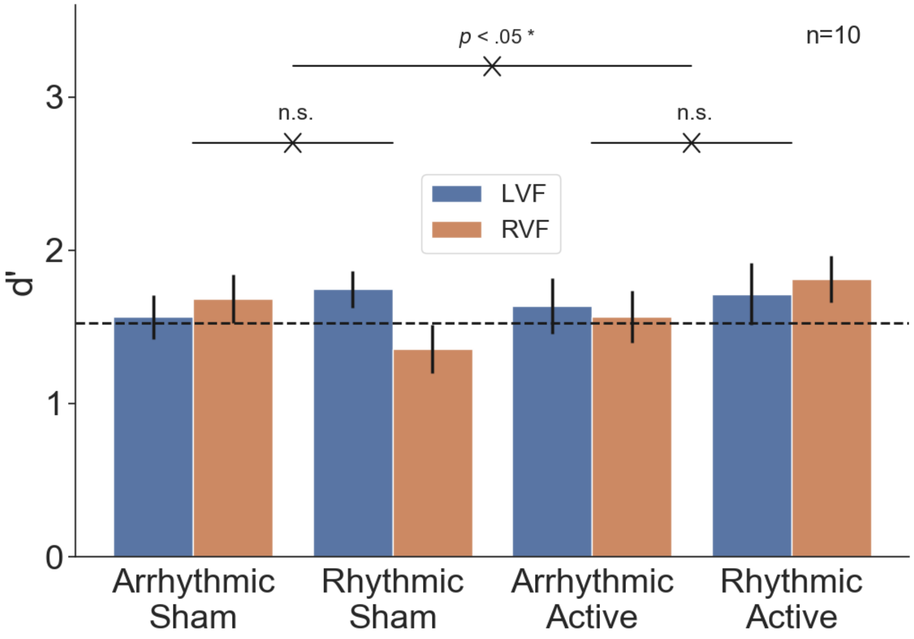


**Figure S2. Sensitivity (d') per condition (rhythmic vs. arrhythmic × active vs. sham stimulation × LVF/RVF target).** The horizontal dashed line indicates the titrated performance level. The 3-way interaction was significant, but the 2-way interactions were not. 10 Hz TMS entrainment of the early visual cortex did not affect sensitivity in the visual discrimination task. The error bars represent ±1 S.E.M. corrected for within-subject design.

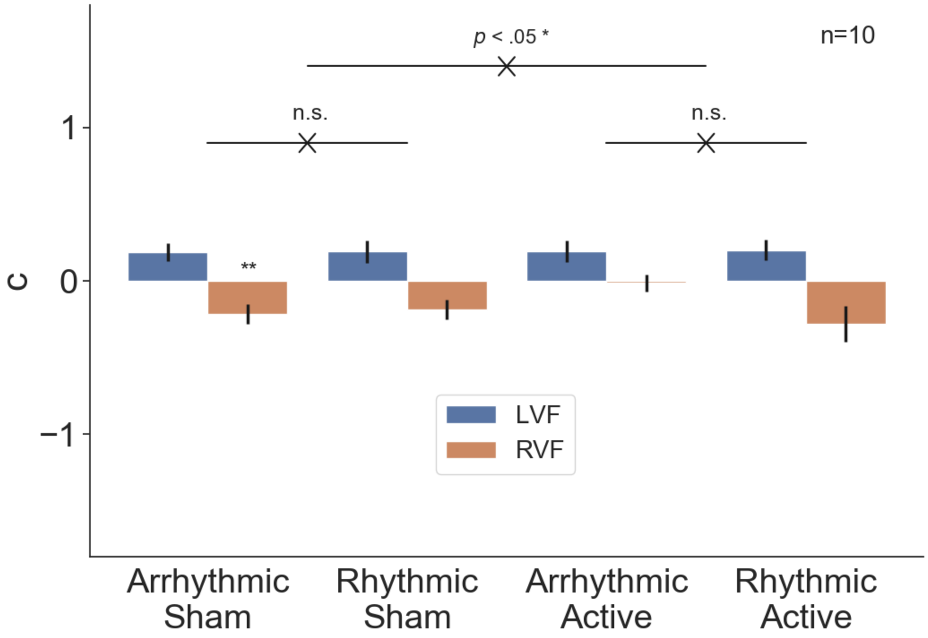


**Figure S3. Criterion (c) per condition (rhythmic vs. arrhythmic × active vs. sham stimulation × LVF/RVF target).** The 3-way interaction was significant, but the 2-way interactions were not. Criterion in all, but one, condition is not significantly different from 0. Overall, 10 Hz TMS entrainment of the early visual cortex did not affect criterion in the visual discrimination task. The error bars represent ±1 S.E.M. corrected for within-subject design.

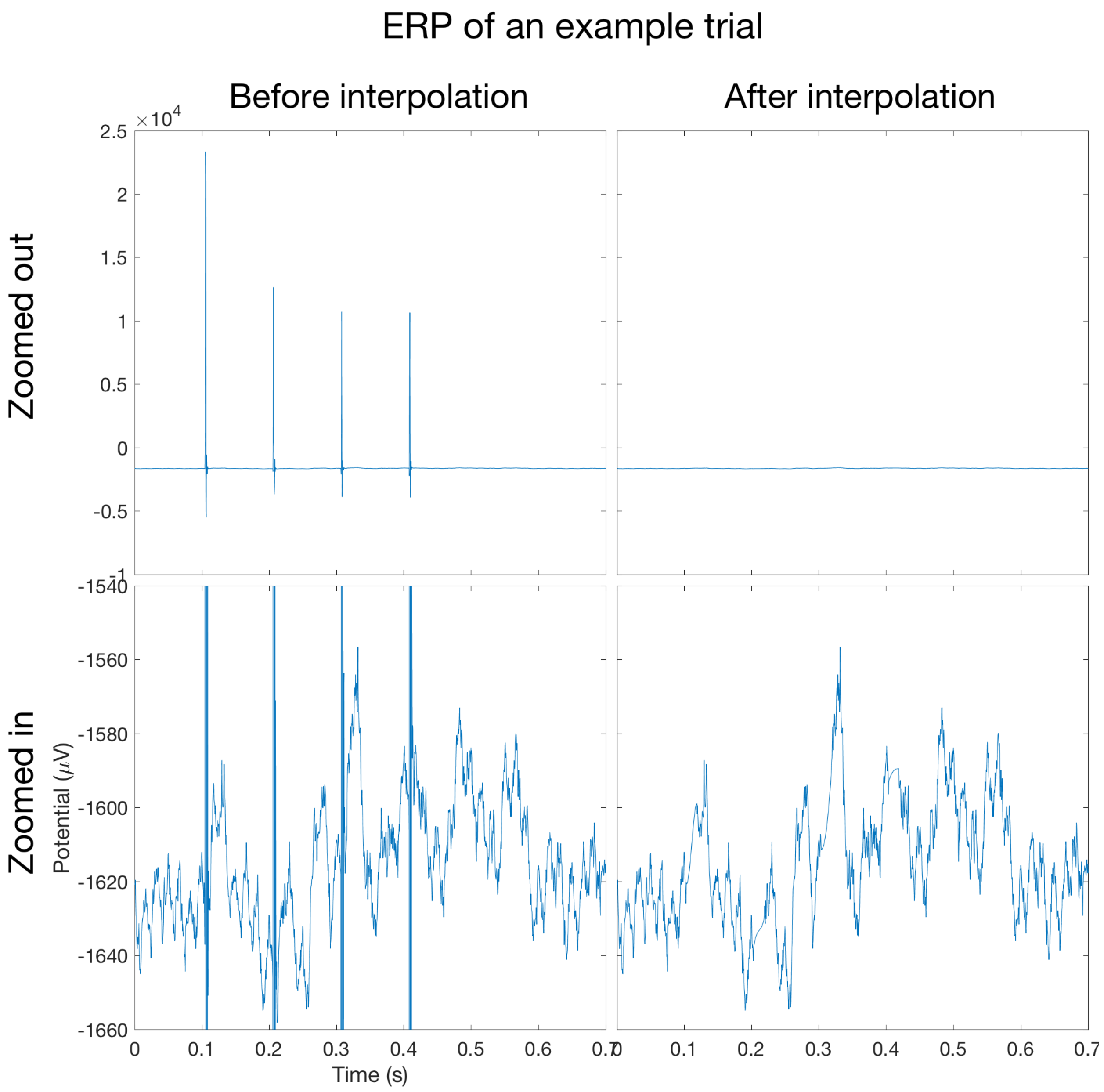


**Figure S4.** TMS spike artifact removal by interpolation in a single trial. All four panels correspond to an example trial in the rhythmic-active condition of participant 03. The top and the bottom rows have different scales for the vertical axis, suitable for the scale of the TMS spike artifact and the actual ERP, respectively. The spline interpolation method reasonably cleaned the EEG signals contaminated by TMS (see Materials and Method). The waveforms reflect the average of electrodes O2 and PO4.

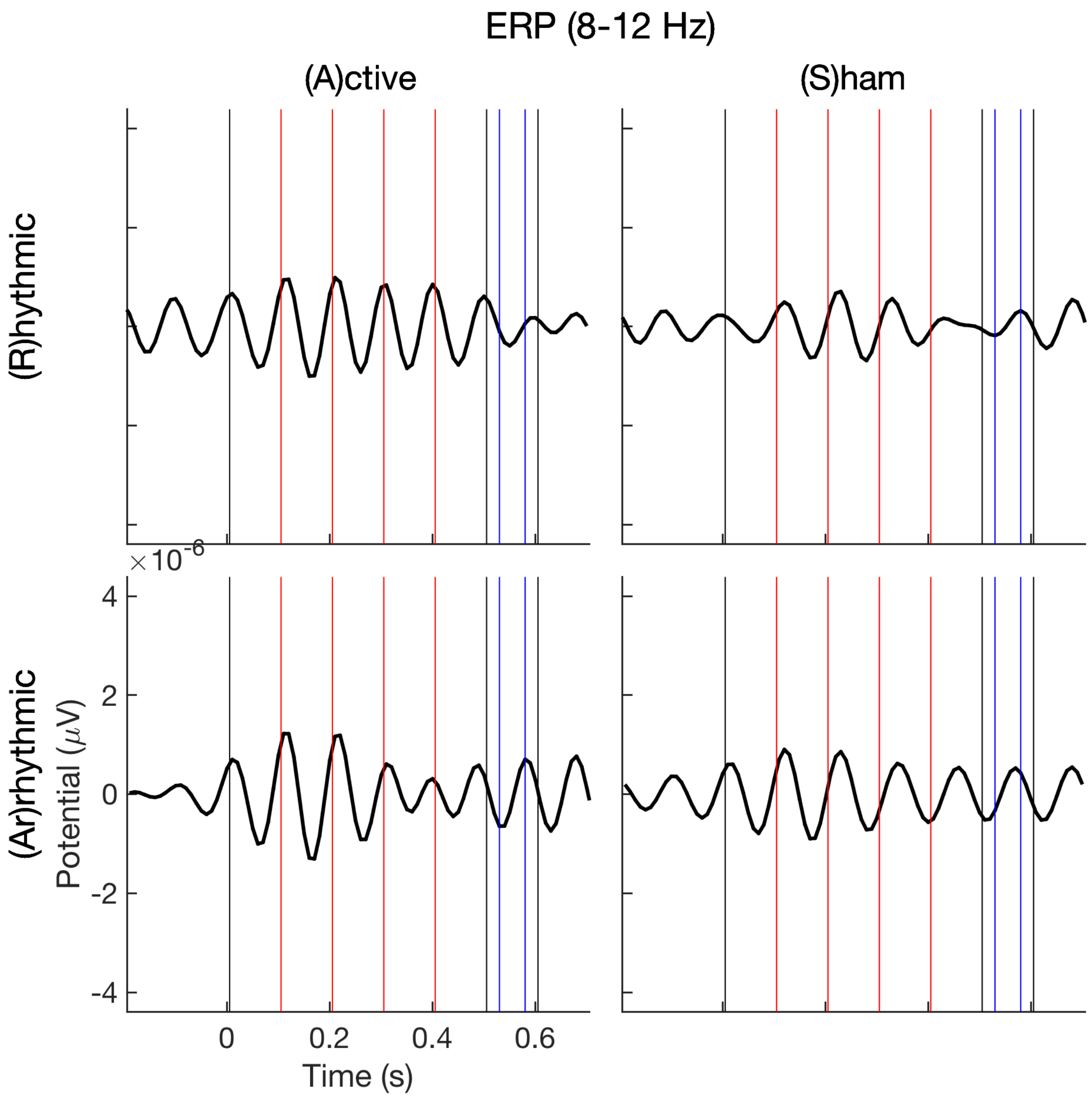


**Figure S5.** Group-averaged ERPs of the four conditions bandpass filtered within the alpha band (8-12 Hz). The forward and reverse Butterworth infinite impulse response filter ensured zero phase shift. The waveforms reflect the average of electrodes O2 and PO4. Each condition is time-locked to the pre-cue onset as t=0. In each panel, the vertical black and red lines demarcate the 100 ms time windows, and the blue lines the Gabor onset and offset. The red lines also indicate the pulse timings in the rhythmic conditions.

**Topographic analysis of phase-locked α-band activities in W0**

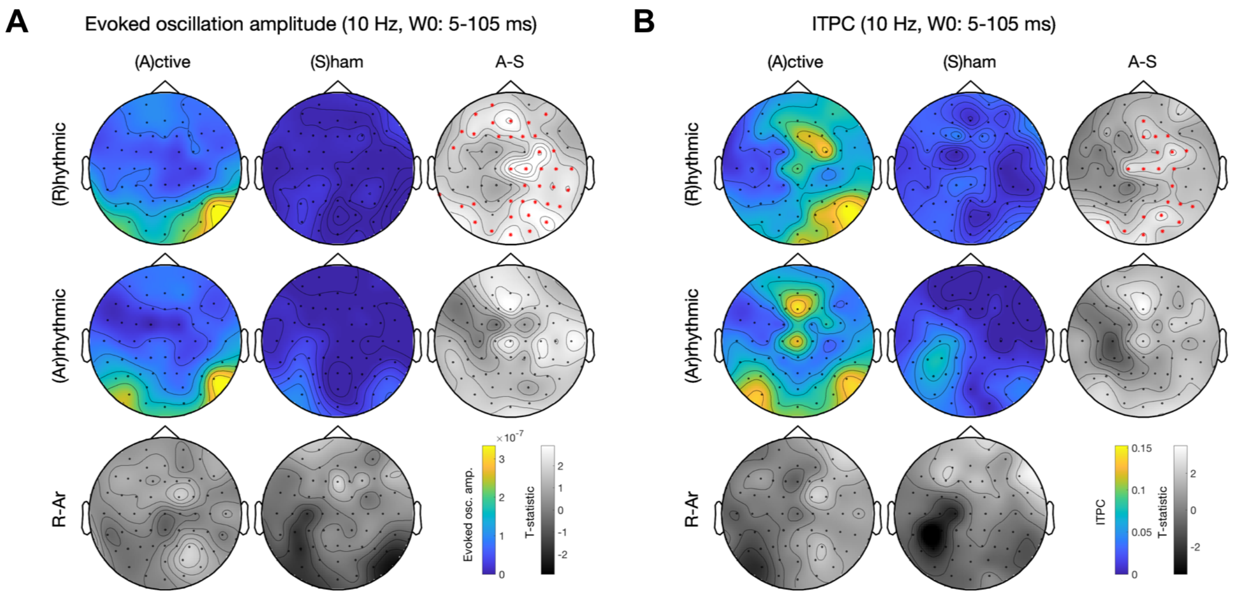


**Figure S6.** Topographic analysis of phase-locked α-band activities in W0. Colored panels are the group-averaged activities per condition (rhythmic vs. arrhythmic × active vs. sham stimulation). Gray panels are the t-statistics of condition contrasts per column or row. The red star symbols indicate significant channels after cluster-based permutation test for multiple comparison correction. (**A**), evoked oscillation amplitude. (**B**), ITPC. Rhythmic-active stimulation elicits widespread activation. The arrhythmic-active stimulation condition is a good control for the rhythmic-active stimulation condition because there is no significant difference before TMS pulses are delivered.

### Topographic analysis of phase-locked α-band activities in W1

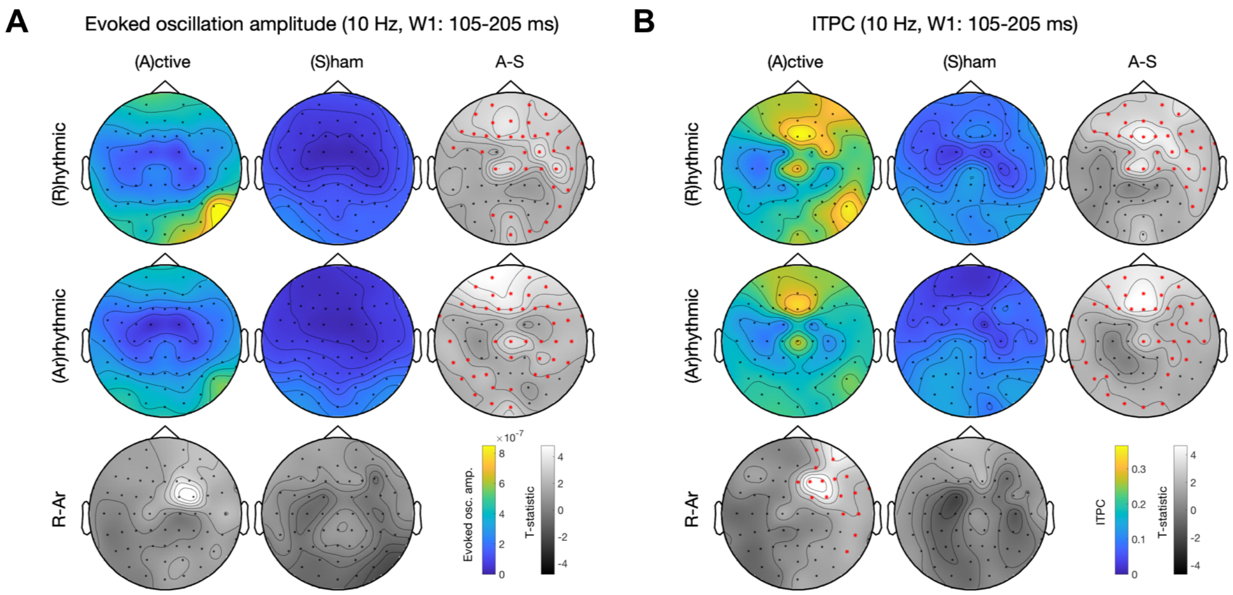


**Figure S7.** Topographic analysis of phase-locked α-band activities in W1. See **Figure S4** for panel layout description. (**A**), evoked oscillation amplitude. (**B**), ITPC. Rhythmic-active stimulation elicits widespread activation. The difference between the rhythmic- and arrhythmic-active stimulation begins in the right frontal region.

#### Topographic analysis of phase-locked α-band activities in W2

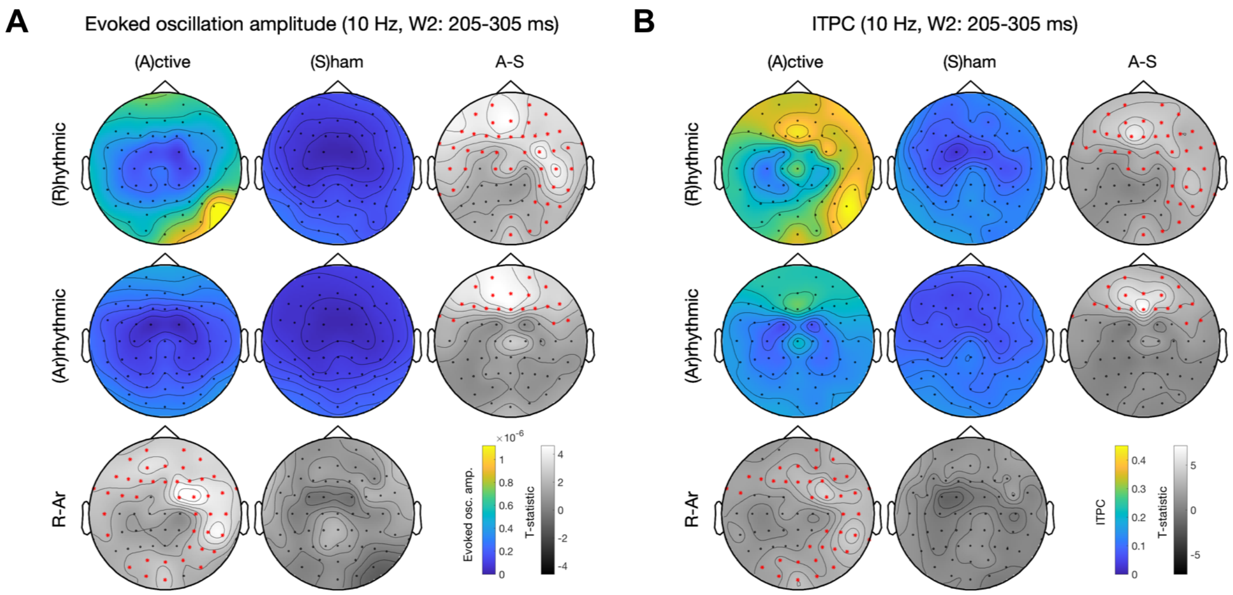


**Figure S8.** Topographic analysis of phase-locked α-band activities in W2. See **Figure S4** for panel layout description. (**A**), evoked oscillation amplitude. (**B**), ITPC. Rhythmic-active stimulation elicits widespread activation. The difference between the rhythmic- and arrhythmic-active stimulation is widespread and starts to include occipital regions.

**Topographic analysis of phase-locked α-band activities in W3**

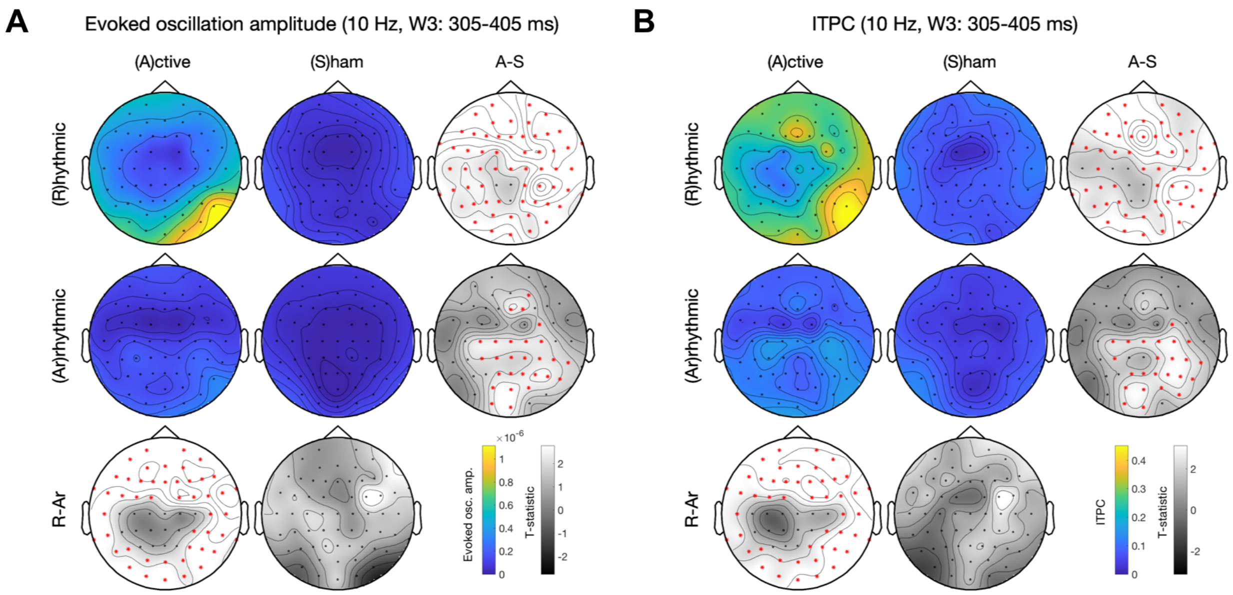


**Figure S9.** Topographic analysis of phase-locked α-band activities in W3. See **Figure S4** for panel layout description. (**A**), evoked oscillation amplitude. (**B**), ITPC. Rhythmic-active stimulation elicits widespread activation. The patterns are similar to those in **Figure S5** and the t-statistics become larger.

**Regression analysis of evoked oscillation amplitude**

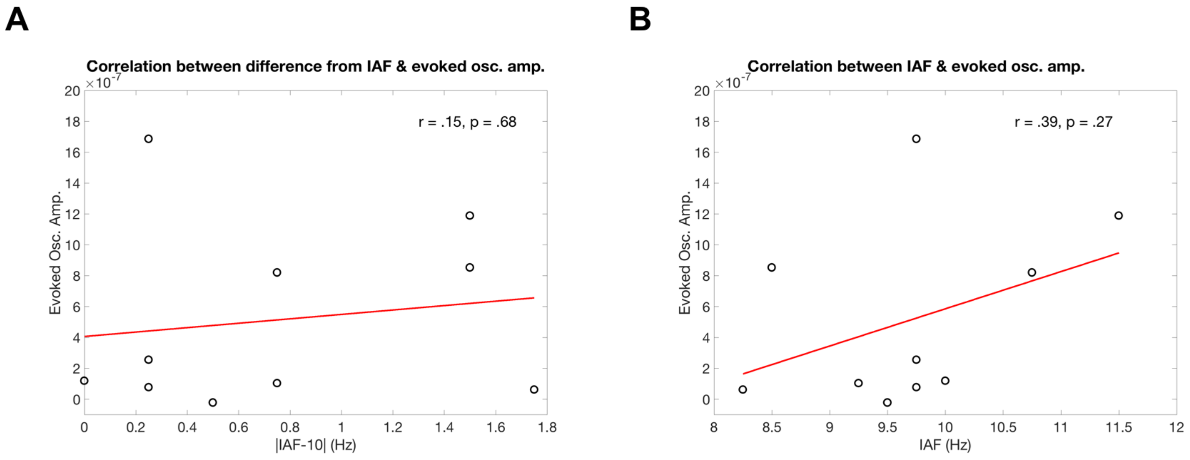


**Figure S10.** Regression analysis of evoked oscillation amplitude in the rhythmic-active stimulation condition. The time window of evoked oscillation amplitude is 0.205-0.705 s, the same as in the significant windows of ITPC in **Table 1**. (**A**), Difference from IAF does not significantly correlate with evoked oscillation amplitude. (**B**), IAF does not significantly correlate with evoked oscillation amplitude.

### Time-frequency analysis of phase-locked activities

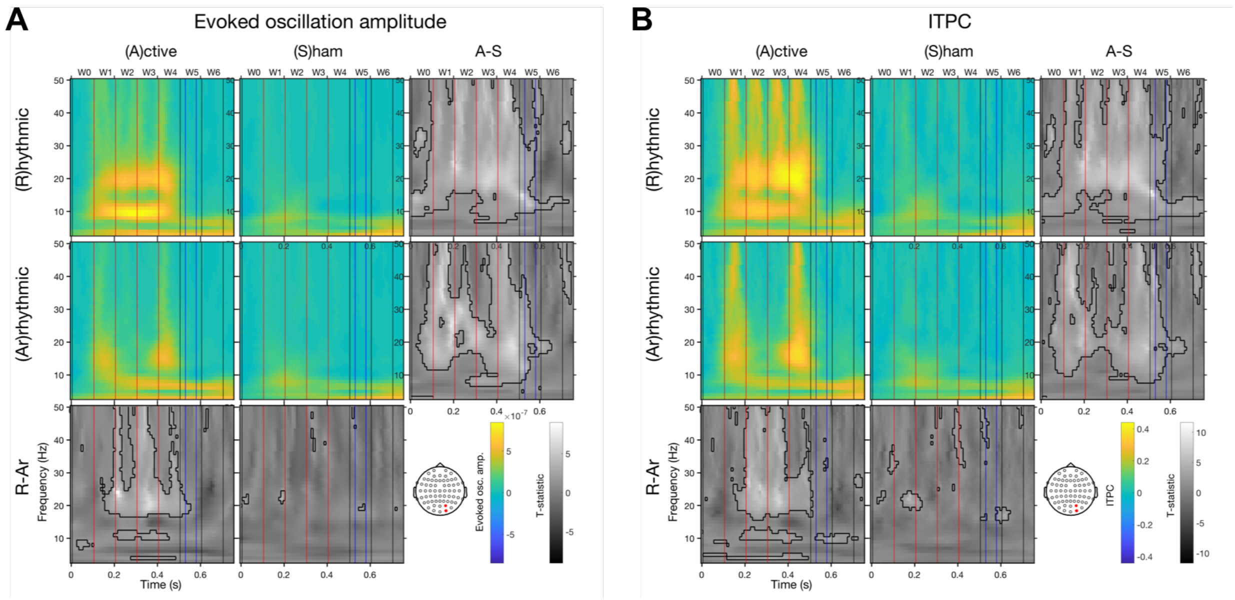


**Figure S11.** Time-frequency analysis of phase-locked activities. The participant whose stimulation site (near POz) deviated from the sites of the rest of participants was excluded from these analyses (n=9). Notice that activity patterns are similar to those presented in **Fig. 2**.

**Regression analysis of ITPC**

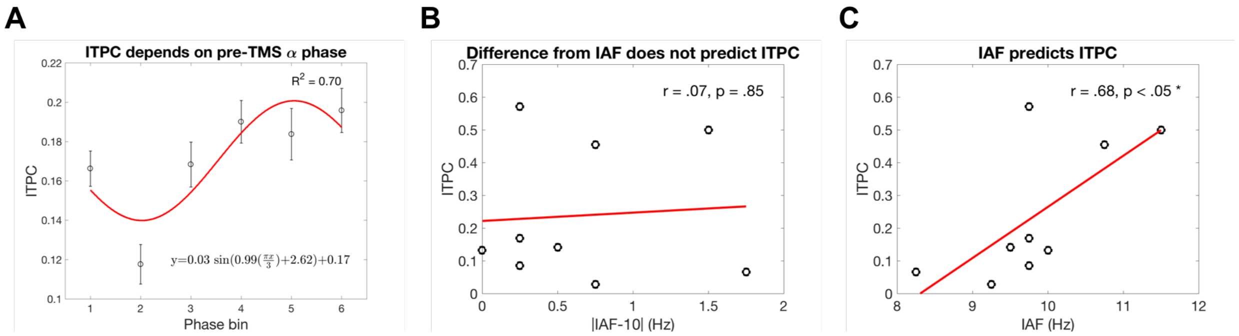


**Figure S12.** Regression analysis of ITPC in the rhythmic-active stimulation condition. The participant whose stimulation site (near POz) deviated from the sites of the rest of participants was excluded (n=9). Notice that activity patterns are similar to those displayed in **Fig. 6**.

**Time-frequency analysis of phase-locked activities**

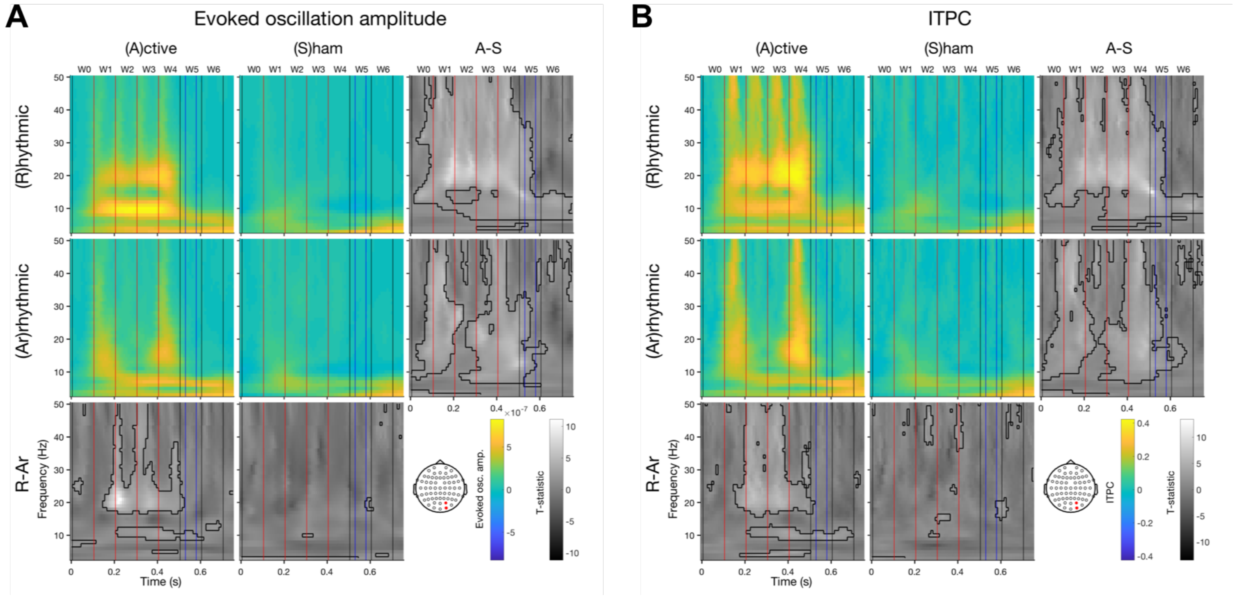


**Figure S13.** Time-frequency analysis of phase-locked activities. Trials following incorrect responses were removed (~24% of total trials). Note that the activity patterns are similar to those shown in **Fig. 2**.

**Regression analysis of ITPC**

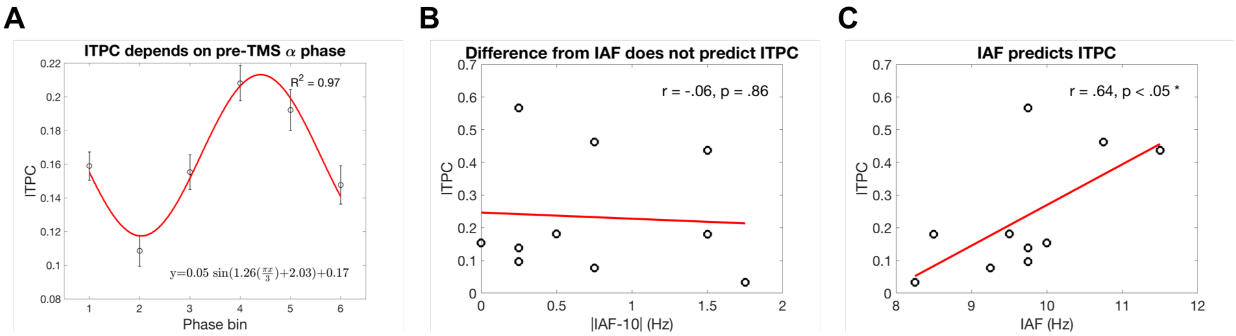


**Figure S14.** Regression analysis of ITPC in the rhythmic-active stimulation condition. Trials following incorrect responses were removed (~24% of total trials). Note that the regression patterns are similar to those shown in **Fig. 6**.

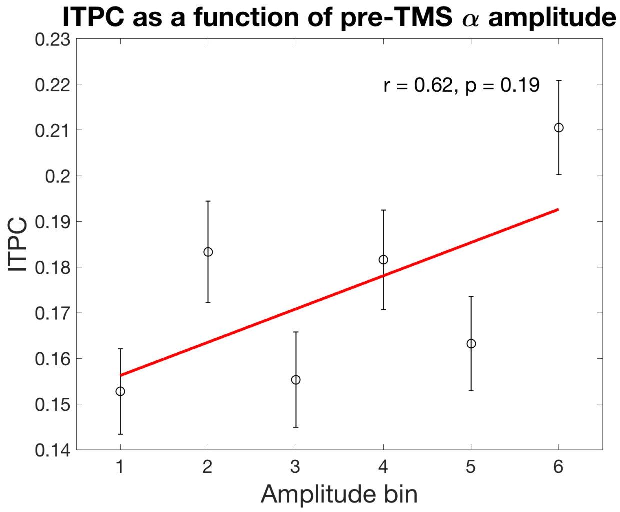


**Figure S15.** Regression analysis of ITPC in the rhythmic-active stimulation condition. The pre-TMS amplitude was sorted into 6 bins. The correlation was not significant.
